# Pre-cancerous Niche Remodelling Dictates Nascent Tumour Survival

**DOI:** 10.1101/2024.07.04.602022

**Authors:** G. Skrupskelyte, J. E. Rojo Arias, Y. Dang, S. Han, M. T. Bejar, B. Colom, J. C. Fowler, P. H. Jones, S. Rulands, B. D. Simons, M. P. Alcolea

## Abstract

Interactions between mutant cells and their environment play a key role in determining cancer susceptibility. However, our understanding of how the pre-cancer microenvironment contributes to early tumorigenesis remains limited. Here, we show that newly emerging tumours at their most incipient stages shape their microenvironment in a critical process that determines their survival. Analysis of nascent squamous tumours in the upper gastrointestinal tract of the mouse reveals that the stress response of early tumour cells instructs the underlying mesenchyme to form a supportive “pre-cancer niche”, which dictates the long-term outcome of epithelial lesions. Stimulated fibroblasts beneath emerging tumours activate a wound healing response that triggers a dramatic remodelling of the underlying extracellular matrix, resulting in the formation of a fibronectin-rich stromal scaffold that promotes tumour growth. Functional heterotypic 3D culture assays and in vivo grafting experiments, combining carcinogen-free healthy epithelium and tumour-derived stroma, demonstrate that the pre-cancerous niche alone is sufficient to confer tumour properties to healthy epithelial cells. We propose a model where both mutations and the stromal response to genetic stress defines the likelihood of early tumours to survive and progress towards more advanced disease stages.

## Background

Groundbreaking studies in human genomics over the past decade have revealed that our healthy tissues accumulate cancer-associated mutations with age^1-5^. These observations highlight new levels of complexity in the early pathophysiology of cancer, questioning what additional factors, beyond cancer mutations, may be playing a role during early carcinogenesis.

Early tumour models spanning a range of epithelial tissues, including oesophagus, skin, and intestine, have started to offer a clearer understanding of what drives tumour initiation^6-11^. Work in this area has shown that tumour formation represents more than the mere accumulation of genetic alterations, highlighting the important role that environmental cues and non-genetic mechanisms, in general, play in this process^8,12-15^. Indeed, mounting evidence indicates that the predisposition of a mutated epithelium to develop tumoral lesions depends on the complex interactions between mutant cells and their dynamic surroundings. Coexisting mutant clones may either synergise or compete, contributing to early tumour initiation^16,17^. And, even after tumours are formed, the presence of neighbouring mutant clones may continue to influence tumorigenesis^6^. Alternative environmental cues such as ECM stiffness^8,13^, as well as direct cell-cell communication between mutant and non-mutant cells^7,9-11,14^, have also been shown to impact mutant clone expansion, susceptibility to tumour initiation, and even invasion^18-20^. Despite this, our understanding of the mechanisms by which environmental factors determine the formation, as well as the long-term survival of emerging tumours, remains limited.

Previous studies using an oesophageal early-tumour model demonstrated that not all nascent tumours have the same chance of survival, with most being cleared from the tissue soon after formation through competition with neighbouring mutant clones^6^. However, what enables the survival of the few remaining tumours and, in particular, how their interaction with the environment during pre-cancer stages shapes their long-term outcome is unknown. We hypothesized that these long-lasting tumours hold the key to understand early tumour survival and offer a window of opportunity to explore the mechanisms modulating early pre-cancer progression. Here, we combine single-cell RNA sequencing with lineage tracing, and 3D heterotypic cultures to study the unique features of the few tumours that manage to escape the pre-neoplastic tumour clearing process in the upper gastrointestinal tract. We demonstrate that, during the most incipient stages of tumour development, fibroblasts react to the pre-neoplastic epithelium by promoting the formation of a fibrotic pre-cancer niche (PCN) that, in turn, feeds back on the epithelium favouring early tumour growth and survival.

### Early tumour survival is linked to the formation of a pre-cancer niche

To study the processes underlying tumour survival at early pre-neoplastic stages, we made use of the DEN (diethyl-nitrosamine) early tumour model, based on a mutagen found in tobacco smoke^6,21^. This well-established carcinogen model recapitulates the mutational landscape of the normal human ageing oesophagus^22^, and induces pre-neoplastic squamous tumours in the upper gastrointestinal mouse tract (GI; including oesophagus and forestomach) (**Fig. 1a, Extended Data Fig. 1a**). Epithelial tumours can be detected in whole tissue whole-mounts from their most incipient stages, i.e. from 10 days post-DEN administration (post-DEN), using keratins KRT6A and KRT17 as markers^6^ (**Extended Data Fig. 1b-e**). Nascent pre-malignant tumours are microscopic, containing as few as 10-20 cells, and are characterised by their distinctive rosette-like structure^6,21^ (**Extended Data Fig. 1d**). This brief window of formation is followed by a tumour clearing process, where the majority of the tumours that initially emerge across the oesophagus are progressively eliminated^6^ (dropping by more than a factor of 3 from 10 days to one year post-DEN; **Extended Data Fig. 1b**). Those that do survive persist largely as hyperplasias or low-grade dysplasias (i.e., pre-neoplastic/pre-cancer tumours stages) but some sporadically grow and progress towards invasive squamous cell carcinomas (**Extended Data Fig. 1g**)^6^, effectively mimicking human cancer progression. As a result of this tumour sorting process, only a subset of the original tumours persists long-term, enabling us to study the mechanisms modulating pre-cancerous tumour survival.

**Fig. 1:**
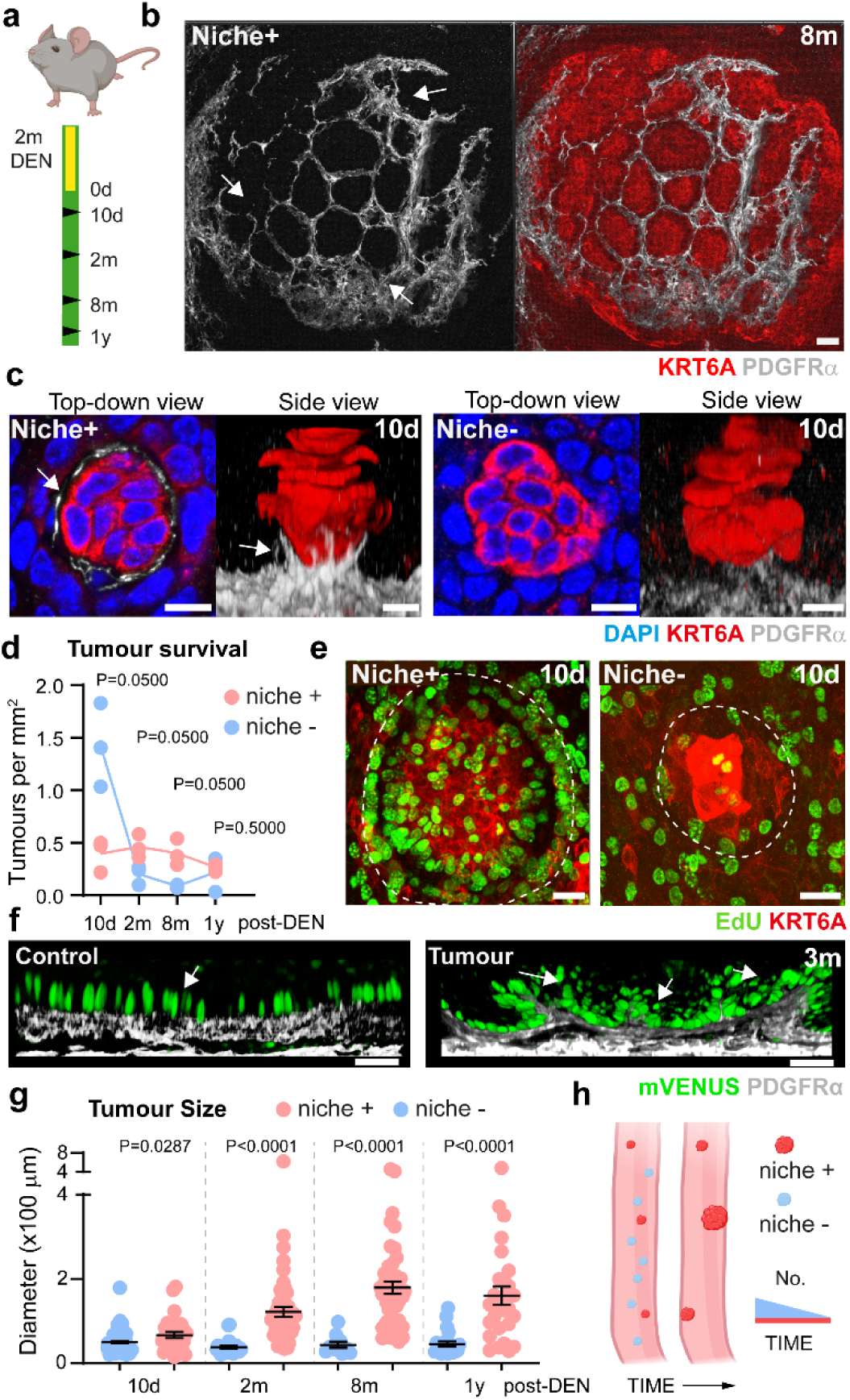
Early tumour niche remodelling is linked to long-term survival. **a**, Experimental DEN (diethyl-nitrosamine) carcinogen protocol. Wild-type mice were exposed to DEN in the drinking water for 2 months. Tissues were collected at the indicated time points. **b, c** Representative confocal images of long-term persisting tumours 8 months post-DEN (**b**) and nascent tumours 10 days post-DEN (c), stained for DAPI (blue), KRT6A (tumour marker, red) and PDGFRα (fibroblast marker, greyscale). Scale bars 25 µm **b**, 10 µm **c**. White arrows mark the remodelled stroma, referred to as niche. **d**, Density of niche+ and niche- tumours at the indicated time points post-DEN treatment. n = 3 mice per time point; solid lines represent means. One-tailed Mann-Whitney test niche*- versus* niche+ tumours. **e**, Confocal images showing EdU incorporation (green) in KRT6A+ nascent tumours (red, dashed line) 10 days post-DEN. Scale bars, 25 µm. **f**, 3D-rendered confocal side-views of a tumour 3 months (m) post-DEN and its respective age-matched control showing R26^Fucci2aR^ tissue (mVenus, S/G2/M cells; green). PDGFRα labels fibroblasts in greyscale. White arrows point to proliferative activity in basal keratinocytes in contact with the stromal. Scale bar, 50 μm. **g,** Diameter (x100 µm) of niche+ (red) and niche- (blue) tumours at the indicated time points. 10-96 tumours were quantified per animal, n=3 mice per time point. Data expressed as mean ±s.e.m. Two-tailed Welch’s t-test niche- *versus* niche+ tumours. **h,** Cartoon illustrating the association between tumour niche remodelling and long-term tumour survival. Red, surviving tumours; blue, vanishing tumours. Panels (**a**) and (**h**) created with BioRender.com.

To identify the processes that may be driving early tumour survival, we first set out to identify the unique phenotypic traits of **long-term persisting tumours** (over 8 months post-DEN) when compared to **nascent tumours** (10 days post-DEN). Histological analysis showed that persisting pre- neoplastic tumours at late time points were characterised by a prominent remodelling of their underlying stroma, forming a nest-like structure composed of stromal fibroblasts (PDGFRα+). Nest fibroblasts spread and protruded into the epithelial compartment, seemingly acting as a supportive scaffold or “**pre-cancer niche**” (PCN; **Fig. 1b, Extended Data Fig. 2a**). Further inspection at early time points revealed that this nest-like stromal structure could also be found in nascent tumours. However, at these early stages, the presence of this structure was largely heterogeneous, with the majority of epithelial lesions (∼70%, 199 lesions out of 296 per oesophagus, on average) showing no apparent stromal reorganization (**Fig. 1c**). This denoted the existence of two phenotypically different nascent tumour types, hereafter referred to as **niche+** and **niche-** (**Fig. 1c**). Histological characterization of the pre-neoplastic stromal scaffold revealed that the major cell component of the nest structure were fibroblasts (**Extended Data Fig. 2b-d**), and showed that remodelled nascent tumours retained an intact basement membrane with no signs of invasion (**Extended Data Fig. 2e**).

Next, we assessed the dynamic nature of the two nascent tumour types. We found that the number of niche+ nascent tumours, despite constituting the minority of all tumours, remained constant over time (**Fig. 1d**). Meanwhile, the number of niche- nascent tumours decreased drastically, effectively accounting for the overall tumour reduction previously observed in this model^6^ (**Fig. 1d; Extended data Fig. 1b**). As a result, the tissue became progressively enriched in niche+ tumours, with the majority (∼82%, 65 out of 79 per oesophagus, on average) showing a supportive stromal scaffold by 8 months post-DEN (**Extended data Fig. 2f**).

The progression towards niche+ tumours prompted us to explore whether stromal remodelling was associated with early tumour survival. Presence/absence analysis across different time points post- carcinogen administration revealed that niche+ lesions were hyperproliferative, and more likely to persist and grow in size compared to tumours lacking this structure (**Fig. 1e-g; Extended data Fig. 2g-j**). Close analysis indicated that keratinocytes in direct contact with the tumour scaffold showed particularly high proliferative activity (**Fig. 1f; Extended data Fig. 2h**), suggesting active communication between the stromal nest and the overlying epithelium. Overall, we observed the tissue gradually becoming enriched in niche+ tumours over time, while niche- lesions were unlikely to persist or grow (**Fig. 1h**). These results were further reinforced by observations in the squamous stomach, where long-term pre-neoplastic tumours also exhibited a remodelled stromal niche (**Extended data Fig. 2k,l**).

Collectively, our data support a model in which the remodelled stromal scaffold acts as a **“pre-cancer niche**” (PCN) promoting tumour growth and survival. These observations link stromal remodelling in nascent tumours with pre-neoplastic tumour progression.

### Pre-cancer niche signals confer tumour properties to mutant-free epithelium

To test the relevance of the pre-cancer niche, we studied the direct impact of the early tumour stroma on healthy epithelial tissue from untreated animals. We anticipated that, if signals from the niche were promoting tumorigenesis, the early tumour stroma may be sufficient to confer tumour features to epithelial cells never exposed to carcinogens. To explore this, we assembled 3D heterotypic cultures in which healthy epithelia from untreated mice were grown *ex vivo* on denuded tumour stroma (lacking the epithelial compartment) from DEN-treated mice. Such an approach allowed us to investigate how pre-existing signals in the early tumour microenvironment affect epithelial cell behaviour^23^ (**Fig. 2a**). By using reporter mouse lines to track the tissue of origin, we were able to reveal that exposure to the tumour stroma by itself drives the emergence of early tumour features in healthy epithelial cells, including a tumour-like morphology (**Extended data Fig. 3a**). Strikingly, healthy untreated epithelial cells exposed to the tumour niche also became highly proliferative, reaching levels similar to those seen in early tumours *in vivo* (**Fig. 2b-d; Extended data Fig. 3b,c**).

**Fig. 2:**
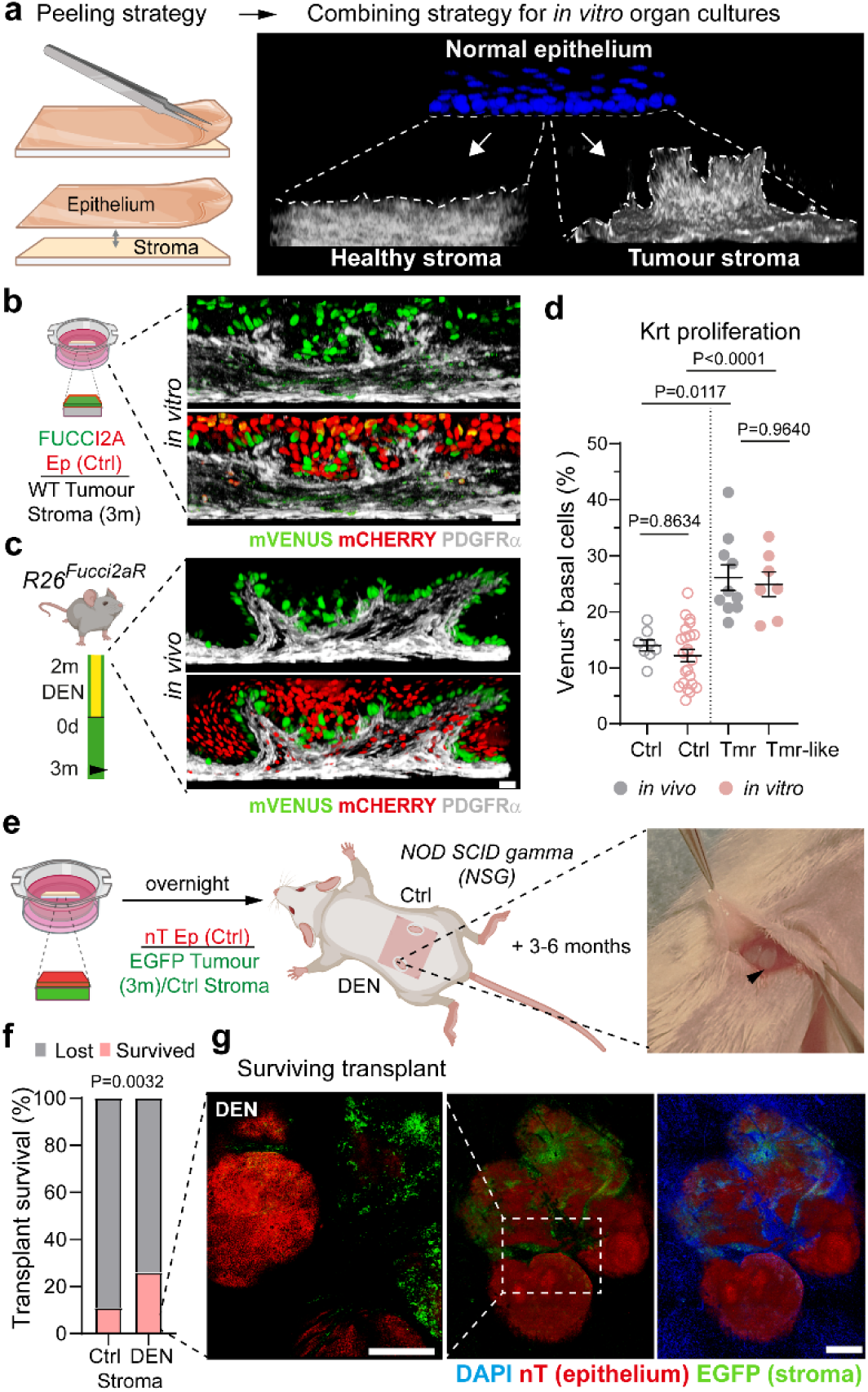
The pre-cancer niche independently modulates keratinocyte proliferation and promotes long-term survival. **a**, Schematic illustrates *ex vivo* tissue recombination approach (3D heterotypic organ culture). After peeling off epithelial and stromal layers, healthy untreated epithelium was combined with stroma of surviving tumours 3 months post-DEN (dashed lines). This was compared to tissue constructs combining healthy untreated epithelium and stroma combined (*in vitro* control). **b-c,** Experimental protocol (left) and representative side-views of 3D-rendered confocal images (right), of 3D heterotypic tissue constructs 7 days post-culture (b) and *in vivo* tumours (c) for comparison. PDGFRα labels fibroblasts (greyscale); in *R26^Fucci2aR^* tissue mCherry, G1 cells; mVenus, S/G2/M cells. Scale bars, 25 µm. Ep, epithelium; wt, wild-type. DEN treated tissue was collected 3 months (m) post-treatment. **d**, Quantification of cycling (mVenus+) basal keratinocytes (Krt) from **b**, and **c** expressed as percentage of total number of basal cells. Dots are data from an individual tumour (Tmr, *in vivo*), tumour-like structure (Tmr-like, heterotypic culture), and the respective *in vivo* and *in vitro* control areas (Ctrl). n=9-23 areas, from 3-6 biological replicates. Mean ± s.e.m whiskers shown.One-way Brown-Forsythe tests and Welch’s ANOVA with multiple comparisons were used. **e**, Schematic representation of the subcutaneous heterotypic grafting approach into NOD SCID gamma (NSG) immunodeficient mice. Grafted epithelium expressed nuclear tdTomato (nT, from nTnG mice), while accompanying stroma was EGFP (from H2B-EGFP mice) to distinguish the origin of cells in the graft. DEN treated tissue was collected 3 months (m) post-treatment. **f**, Quantification of surviving and lost graft constructs from **e** 3-6 months post-transplantation expressed as percentage. 19 control (Ctrl) and 23 DEN constructs were transplanted across 10 animals, respectively. Statistical significance was determined by a one-sided Chi-squared test. **g,** Representative confocal image (from **f**) showing the long-term survival of a graft combining healthy untreated nT epithelium with EGFP+ tumour stroma (scale bars 500 µm). Panels (**a**), (**b**) and (**e**) created with BioRender.com.

Follow-up experiments investigated whether the tumour niche pro-survival phenotype was also observed *in vivo*. Heterotypic tissue constructs grafted into immunodeficient mice revealed that healthy epithelial cells exposed to early tumour stromal signals were more likely to engraft long-term than those that received signals from control stroma from untreated animals (**Fig. 2e-g; Extended data Fig. 3d**), supporting the pro-survival effect of the early tumour niche.

Overall, our observations suggest that, alongside epithelial mutations, the early tumour microenvironment, and in particular the mesenchymal tumour niche, is able to promote epithelial cell growth, favouring pre-cancerous tumour survival and, ultimately, disease progression.

### Tumour cell heterogeneity is associated with early tumour stromal remodelling

In order to identify the mechanisms leading to the formation of the pre-cancerous niche, we set out to study the communication between tumour cells and their underlying mesenchyme in surviving tumours. To this end, we performed single-cell RNA sequencing (scRNA-seq) analysis of the squamous upper GI tract 8-months post-DEN administration (**Fig. 3a**). At this strategically chosen time point, the tissue is enriched in enlarged niche+ surviving tumours at pre-neoplastic stages, i.e., dysplasias (**Fig. 1b,d,g; Extended Data Fig. 1b,g; Extended Data Fig. 2f**). Given the microscopic size of the pre-neoplastic tumours, we devised a punch biopsy strategy to enrich our scRNA-seq sample preparations in tumour cells. Individual tumours were cut out using submillimetric biopsy punches (as small as 0.5mm in diameter) and pooled prior to tissue dissociation (**Fig. 3a**; **Extended Data Fig. 4a,b**). The transcriptional signature of pre-neoplastic tumour cells (**tumour**) was then compared to that of cells from tumour-free areas from DEN-treated mice (**DEN**; internal control), as well as control tissue cells from healthy untreated animals (**Ctrl**; **Extended Data Fig. 4c,d**). Punches contained both epithelial and stromal cell populations (**Fig. 3b, Extended Data Fig. 4e,f,g**). To identify the relevant cell populations/states involved in the epithelial-mesenchymal cross-talk, we investigated the transcriptional signature of these two cell compartments (epithelial and mesenchymal) separately.

**Fig. 3:**
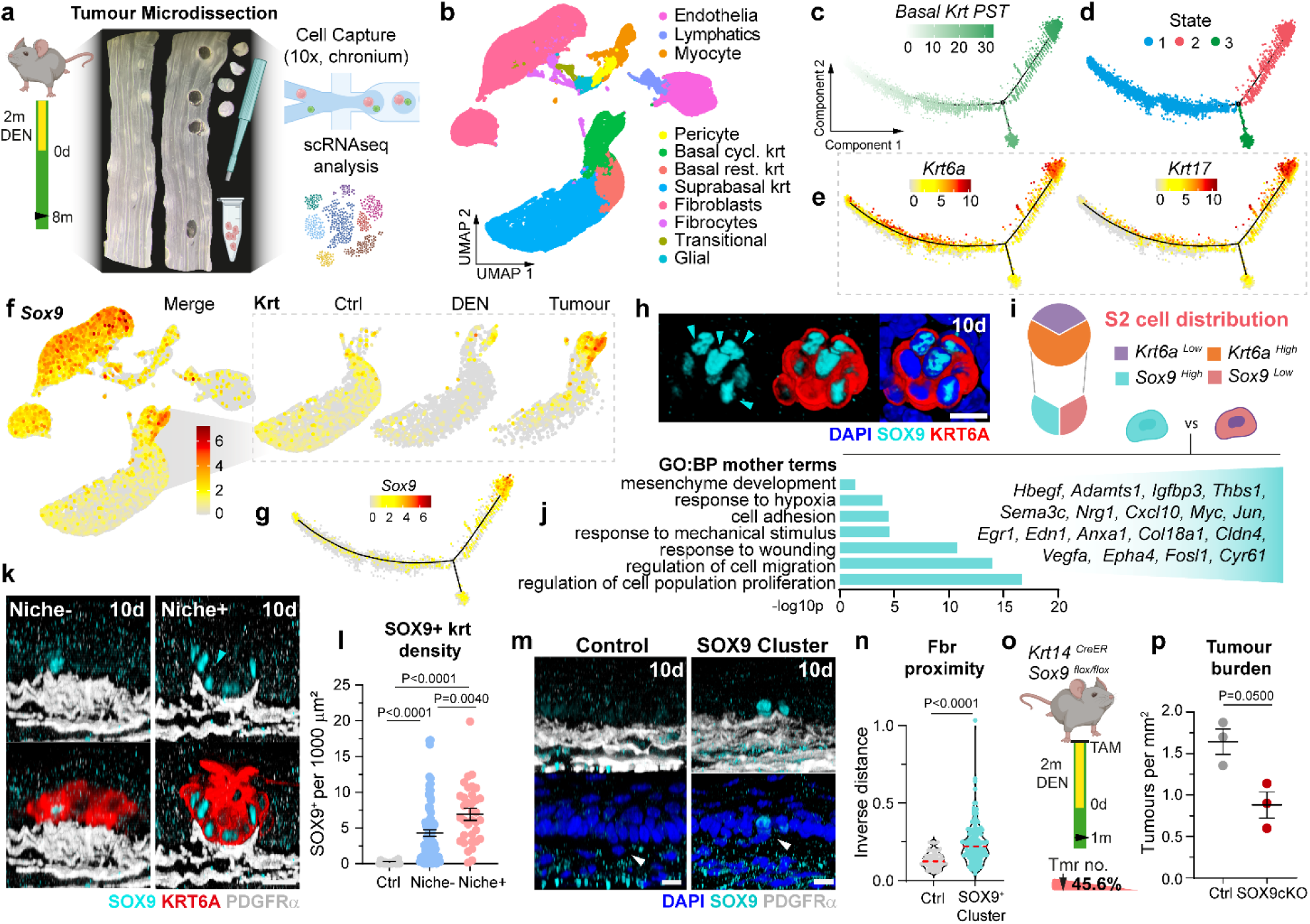
SOX9^high^ epithelial cells are associated with early tumour stromal remodelling. **a**, Tumours from the squamous upper gastrointestinal tract were micro-dissected using a punch biopsy approach (0.5-1mm diameter biopsies). Tumour samples, equivalent non-tumour areas in DEN treated mice (DEN), as well as healthy untreated controls (control) were dissociated and subjected to single-cell RNA sequencing (scRNA-seq). Tumours were collected 8 months (m) post-DEN, or equivalent age-matched controls. **b**, UMAP showing the cell type annotation. Cycl, cycling; rest, resting; krt, keratinocytes. **c**, Basal keratinocyte two-dimensional pseudotime (PST) trajectory, and respective pseudo-stable cellular states S1-S3 (**d**). **e,** Expression of tumour makers *Krt6a* and *Krt17* in PST trajectory. **f, g** UMAP projection and PST trajectory showing expression of *Sox9* expression. Insets in **f** depict the overexpression of *Sox9* gene in tumour-derived keratinocytes. **h**, Representative confocal images showing heterogeneous SOX9 expression (cyan) among KRT6A^high^ cells (red) in nascent tumours (10d post-DEN). Cyan arrowheads set SOX9^high^ cells apart from SOX9^low^ cells. Scale bars, 10 µm. **i**, Pie chart depicting the distribution of *Krt6a* and *Sox9* expressing cells in PST State-2. **j,** Gene Enrichment Analysis denotes the different transcriptional signature of the two tumour cell states identified, *Krt6a*^high^/*SOX9*^low^ and *Krt6a*^high^/*SOX9*^high^. Terms ranked by p-value; representative genes listed on the right. **k**, Representative images showing SOX9 distribution in niche- and niche+ nascent tumours (10 days post-DEN). Cyan arrowheads highlight SOX9 and stroma proximity. DAPI, blue; SOX9, cyan; KRT6A, red; PDGFRα, greyscale. Scale bars, 10 µm. **l**, Quantification of SOX9^high^ cells in **k** normalised per area. Ctrl, equivalent areas in aged-matched healthy untreated tissue. n=35-84 tumours and n=19 Ctrl areas, 3 mice per condition. One-way Welch ANOVA comparing tumour types and control. **m**, Confocal images of *Krt6a*^high^/*SOX9*^high^ clusters in tumour-free areas of DEN-treated epithelium 10 days post-DEN. DAPI, blue; SOX9, cyan; PDGFRα, greyscale. Scale bars, 10 µm. White arrowheads highlight keratinocyte to fibroblast proximity. **n**, Quantification of fibroblast proximity to basal layer from **m,** expressed as the inverse of the distance between basal keratinocytes and underlying fibroblasts. n=28-33 clusters or equivalent control areas from 4 mice. Two-tailed Mann-Whitney test comparing the regions. **o**, Experimental protocol: *Krt14^CreER^Sox9^flox/flox^* mice received a dose of Tamoxifen (TAM) followed by the DEN. Tissues were collected 1- month post-DEN and compared to DEN-treated un-induced controls. Tumour burden decreased ∼45.6% in *Krt14^CreER^Sox9^flox/flox^*induced mice (SOX9cKO). **s**, Quantification from **o**. n = 3 mice per time point; data is presented as mean±s.e.m. One-tailed Mann-Whitney test. Panels (**a**) and (**o**) created with BioRender.com.

We started by exploring the epithelial transition from healthy to pre-neoplastic states by pseudotime analysis of the basal cell population, which comprises progenitor cells. The analysis revealed distinct states (**Fig. 3c,d; Extended Data Fig. 5a**). States 1 and 3 included cells from all conditions (Ctrl, DEN and tumour) and showed the typical signature of basal epithelial cells, as previously reported^24^. In particular, state 1 comprised basal cells committed to differentiation, while state 3 included basal proliferating cells (**Extended Data Fig. 5b-e**). State 2, instead, represented a distinctive tumour state, with cells expressing high levels of *Krt6a* and *Krt17* (**Fig. 3e; Extended Data Fig. 5f**), two established early tumour markers in the DEN carcinogen model^6,21^ (**Extended Data Fig. 6a**). The hyperproliferative phenotype (**Extended Data Fig. 5b**) of state 2 cells was marked by a downregulation of the TGFβ signalling pathway (*Smad7, Id1, Id2, Id3, Id4, Bambi, Tgif*)^25^, which has previously been linked to rapidly proliferating keratinocytes^26^ (**Extended Data Fig. 5f)**. The tumoral nature of state 2 cells was further supported by gene enrichment analysis, which showed an increased expression of genes related to cancer associated processes such as hypoxia (*Hif1a*, *Vegfa*, *Edn1*, *Cited2*, *Sirt1*), cell adhesion (*Sdc4, Clu, Cyr61, Ephb3, Thbs1*), migration (*Plet1, Actg1, Myadm, Krt16*), as well as to stress-induced pathways such as EGFR (*Ereg*, *Epgn*, *Areg*, *Hbegf, Egr1, Lamc2*^27^), p53, IL-17, TNF and NFκB (*Gadd45a*, *Igfbp3, Tnfaip3*, *Nfkbia*, *Sesn2, Cxcl10, Cxcl2, Ccl20, Ptgs2*) **(Extended Data Fig. 5f)**. This was further reinforced by increased levels of transcription factor (TF) genes associated with a tumour stress response (*Sox9, Jun, Fos, Atf3, Egr1*; **Fig. 3f,g**; **Extended Data Fig. 6b,c**)^28-32^.

One gene, *Sox9*, was of particular interest, given its role in tissue stromal remodelling during morphogenesis, epithelial branching and cancer invasion, as well as its involvement in fate plasticity and early tumorigenesis^28-30,33,34^. Whole-mount image analysis confirmed that, unlike healthy control tissue, nascent tumours (10 days post-DEN) express SOX9 (**Fig. 3h**), albeit showing a largely heterogeneous profile. That is, early tumours showed marked differences in the proportion of SOX9- expressing cells **(Extended Data Fig. 6d)**. Differential gene expression analysis revealed pronounced differences in the transcriptional profile of *Krt6a*^high^/*Sox9*^high^ and *Krt6a*^high^/*Sox9*^low^ (hereafter referred as *Sox9*^high^ and *Sox9*^low^) tumour cells, thereby defining two distinct cell states within the pre-neoplastic tumour (**Fig. 3i,j; Extended Data Fig. 6e**). The expression profile of *Sox9*^high^ cells showed a particular enrichment in TFs associated with stress response (*Jun, Jund, Fos, Myc, Epgn, Atf3*)^30,31^, as described in state 2. Additional expression changes included genes known to be associated with stromal communication, including cell adhesion to extracellular matrix (ECM) and cell-to-cell adhesion (*Col18a1, Sema3c, Itga6b, Tnc, Anxa1, Epha4, Edn1,*), as well as ECM break- down (*Adamst1*^35^). Overexpression of cell adhesion genes *Cyr61*^36,37^ and *Thbs1*^38,39^ was of particular interest due to their recognised role in communication with fibroblasts (**Fig. 3j; Extended Data Fig. 6f-h**). Altogether, the data suggest a close communication between *Sox9^high^* epithelial and stromal cells.

Analysis of SOX9 expression in 10 day post-DEN tumours revealed that the proportion of SOX9+ cells, as well as SOX9 expression levels, were both significantly higher in niche+ tumours (**Fig. 3k,l; Extended Data Fig. 6i,j**). Closer inspection showed that SOX9 was expressed primarily in basal cells of early tumours, which are those in direct contact with the niche (**Fig. 3k; Extended Data Fig. 6k**), indicative of an intimate relationship between *Sox9*^high^ cells and the underlying stroma. Overall, these data support a direct link between SOX9 expression and the presence of the pre-neoplastic tumour niche.

To explore whether the SOX9^high^ tumour cell state is associated with the remodelling of the tumour stroma and the formation of the pre-neoplastic tumour niche, we took advantage of sporadic clusters of KRT6A+/SOX9+ cells found in phenotypically normal (non-tumour) areas of DEN-treated tissue, likely marking prospective tumour cells prior to lesion formation. Isolated SOX9+ cell clusters showed signs of fibroblast attraction, presenting an increased density of fibroblasts directly underneath them (**Fig. 3m; Extended Data Fig. 6l,m**). This notion was further reinforced by the closer proximity between SOX9+ keratinocytes and fibroblasts, in comparison to the surrounding SOX9- epithelium (**Fig. 3n; Extended Data Fig. 6n)**.

Overall, this data establishes the KRT6A^high^/SOX9^high^ tumour state as a relevant player in epithelial-stromal communication from the most incipient stages of tumorigenesis, and unveils its potential role in stromal remodelling and early tumour niche formation. Accordingly, SOX9 depletion in Krt14- Cre^ER^; Sox9^flox/flox^ DEN treated mice (**Extended Data Fig. 6o,p**) led to a significant reduction in tumour burden (**Fig. 3o,p**).

### Stromal reorganisation is driven by resident PDGFRα^low^ pro-fibrotic fibroblasts

After identifying *Krt6a*^high^/*Sox9*^high^ as a key epithelial state driving epithelial-stromal cross-talk and early tumour survival, we next turned our attention to the stromal compartment with the aim to identify the fibroblast population involved in early tumour niche formation. Fibroblast scRNA-seq analysis showed a degree of heterogeneity in the expression of the fibroblast marker *Pdgfra*^40^, both in control and tumour samples (**Fig. 4a, Extended Data Fig. 7a**). In line with observations in other epithelial tissues^41^, immunofluorescence analysis revealed a visible compartmentalization of *PDGFR*α^high^ and *PDGFR*α^low^ cells within the tissue stroma, marking the presence of two distinct populations (**Extended data Fig. 7b,c**). PDGFRα^low^ fibroblasts were observed immediately underneath the epithelium, forming the lamina propria (**Extended data Fig. 7c**), a thin loose connective tissue separated from the lower dense irregular submucosa compartment by a single layer of smooth muscle cells (the muscularis mucosa; **Extended data Fig. 7d**)^42^. In contrast, the submucosa, the deepest stromal layer, was populated by PDGFRα^high^ fibroblasts (**Extended data Fig. 7c,d**). Notably, differential gene expression showed that *Pdgfra*^low^ fibroblasts express higher levels of genes encoding for structural (interstitial /scaffolding matrix) ECM components (such as *Fn1, Has2, Col1a1, Col3a1*)^43^, while *Pdgfra^high^* fibroblasts show enrichment in genes encoding for ECM tethering (pericellular/underlying matrix), collagens (such as *Col4a, Col6a*) and other basement membrane components (such as *Lama1, Thbs1*; **Extended data Fig. 7e,f**)^44^.

**Fig. 4:**
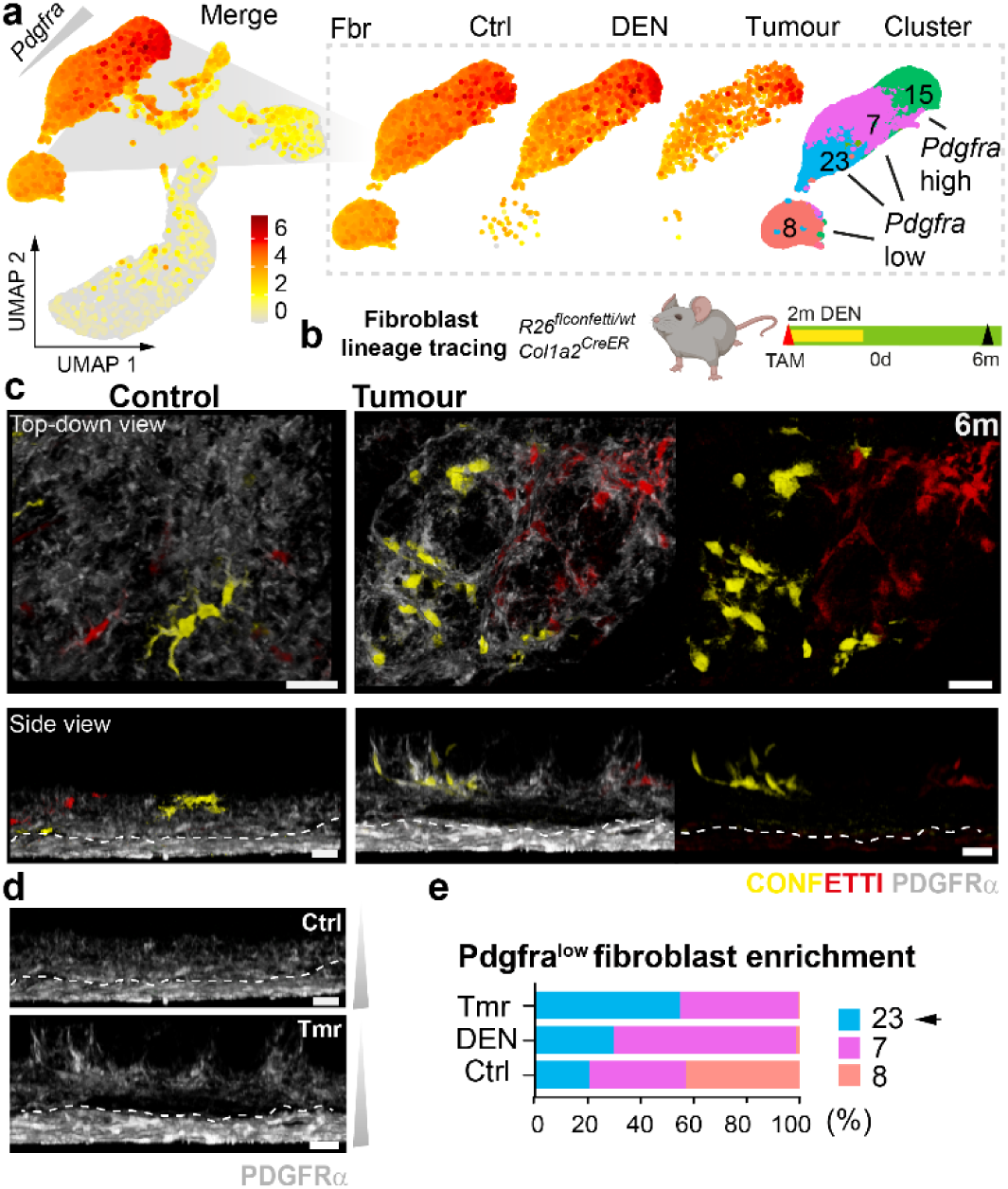
The early tumour niche is formed by an expanded PDGFRα^low^ fibroblast population. **a**, UMAP projection denoting the heterogenous expression of *Pdgfra* in the fibroblast compartment; insets showing expression split per condition: healthy untreated control (Ctrl), non-tumour control in DEN treated mice (DEN), and tumours. Clusters included in the fibroblast analysis are shown (right). **b**, Experimental protocol for fibroblast lineage tracing: *Col1a2^CreER^R26^FlConfetti/wt^* mice received a dose of Tamoxifen (TAM) followed by DEN treatment. Samples were collected 6 months (m) post-DEN treatment. **c-d**, Representative top-down (top) and side-views (bottom) of control or tumour tissue from **b**. PDGFRα, greyscale; lineage traced confetti+ cells (yellow and red). Scale bars, 50 µm. Dashed lines label the transition between two distinct stromal compartments. **d**, Split channel from **c** shows marked difference in PDGFRα expression across these two stromal compartments both in control and Tumour samples; lamina propia, PDGFRα^low^ (underlying the epithelium); submucosa, PDGFRα^high^ (deeper stromal layer). **e**, Stacked barplot showing fibroblast cluster enrichment across conditions. Black arrow marks cluster 23 (from **a**) as a PDGFRα^low^ preferentially enriched in tumours. Panel (**b**) created with BioRender.com.

Given the close proximity of PDGFRα^low^ fibroblasts to the epithelium, we reasoned that this may represent the main population giving rise to the early tumour “niche”. To establish whether this was the case, we used a genetic lineage tracing approach. Col1a2-Cre^ER^; R26^FlConfetti/WT^ mice were treated with Tamoxifen to induce Cre-driven recombination, labelling of fibroblasts both in the lamina propria and the submucosa (**Extended data Fig. 7g-j**). To trace the origin of the tumour “niche”, the induction was performed prior to tumour formation (i.e., prior to DEN administration) **(Fig. 4b).** Clonal analysis 6 months post-DEN administration showed that the contribution of fibroblasts to tumour niche formation was associated with their clonal expansion over time, primarily within the inter-lobe space of the tumour (i.e. within the niche space; **Fig. 4c; Extended data Fig. 7k-p**). Immunostaining confirmed that the clones forming the niche consist of PDGFRα^low^ fibroblasts localised immediately underneath the epithelium (lamina propria; above the dash line **Fig. 4d**), identifying the PDGFRα^low^ population as being of primary interest to further understand the role of epithelial-fibroblast communication during early tumorigenesis.

Next, we interrogated the transcriptional signature of PDGFRα^low^ niche-associated fibroblasts. For this we focused on cluster 23 (C23) fibroblasts, a *Pdgfra*^low^ cluster enriched in tumour cells (**Fig. 4e**). Compared to the rest of *Pdgfra^low^*fibroblasts (C7,C8 and C38), C23 fibroblasts showed an enrichment in genes associated with wound healing/fibrosis (*Fn1, Fbn1, Mfap5, Pdpn, Adam15, Egfr,*)^9,45^ (**Extended data Fig. 8a**). Other genes included *Mfap5*, *Col1a2*, *Col3a1*, previously found to also be overexpressed in cancer-associated fibroblasts (CAFs) in oesophageal cancers^46^. This hinted at the activation of a tissue repair response in the stroma of early tumours, in line with the long-standing notion that tumours are wounds that do not heal^47^. We next assessed whether the niche- forming fibroblasts recapitulated CAF features by comparing the gene expression profile of C23 tumour cells to DEN and Ctrl conditions. Despite a subtle upregulation of CAF-associated genes^12,46,48^, including *S100a4, Vim, Fap, Postn* and *Ctgf*, their protein levels (VIM, FAP, FSP1) remained largely unaltered or non-detectable in the niche of surviving tumours (**Extended data Fig. 8b-e**). Similarly, we did not observe the typical pro-tumorigenic increase in fibroblast proliferation^49^ (**Extended data Fig. 8f**), suggesting that niche+ fibroblasts do not present a fully established CAF phenotype at incipient tumour stages. However, niche+ fibroblasts from nascent tumours did express nuclear YAP (**Extended data Fig. 8g**), indicating the presence of pre-CAFs in the niche, i.e., fibroblasts in a transition state prior to becoming CAFs^50^

To assess whether genetic alterations in the tumour stroma may be driving the tumour niche phenotype, we performed deep-targeted sequencing of 192 cancer-related genes (median on-target coverage 236×)^22^. The data showed minimal mutational burden in the tumour stroma, matching the level of untreated or internal control samples (**Extended data Fig. 8h,i**). The epithelial tumour compartment, instead, served as a positive control presenting a significant level of mutations^22^. These results indicate that DEN administration induces gene perturbations primarily in the epithelial compartment and argues against somatic mutations in fibroblasts being responsible for pre-cancer niche formation.

These observations reveal that changes in the early tumour stroma, prior to the emergence of a CAF phenotype, contribute to the formation of the pre-cancer tumour niche and consequently support tumour survival.

### Epithelial-mesenchymal communication in early surviving tumours

To study epithelial-mesenchymal communication in surviving tumours, we set to identify ligand-receptor interactions^51^ enriched across the different cell populations that co-exist in the DEN-treated tissue (**Fig. 5a,b; Extended data Fig. 9a**). Our analysis included interactions between non-tumour cells, i.e., **control keratinocytes** (*Krt6a*^low^) and **control fibroblasts** (*Pdgfra*^low^; C23), as well as between tumour-associated cells, i.e., **tumour keratinocytes** (*Krt6a*^high^/*Sox9*^low^), **tumour niche keratinocytes** (*Krt6a*^high^/*Sox9*^high^; **Fig. 3h,i**), and **tumour niche fibroblasts** (*Pdgfra*^low^ ; C23; **Fig. 4e**). We selected the top ten outgoing interactions from both keratinocytes and fibroblasts, respectively, and visualised the matching incoming interactions to feature the receiver cells (**Fig. 5c,d; Extended Data Fig. 9b**). We then shortlisted the pathways that predicted stronger communication between tumour niche keratinocytes and tumour niche fibroblasts (**Fig. 5e,f**).

**Fig. 5:**
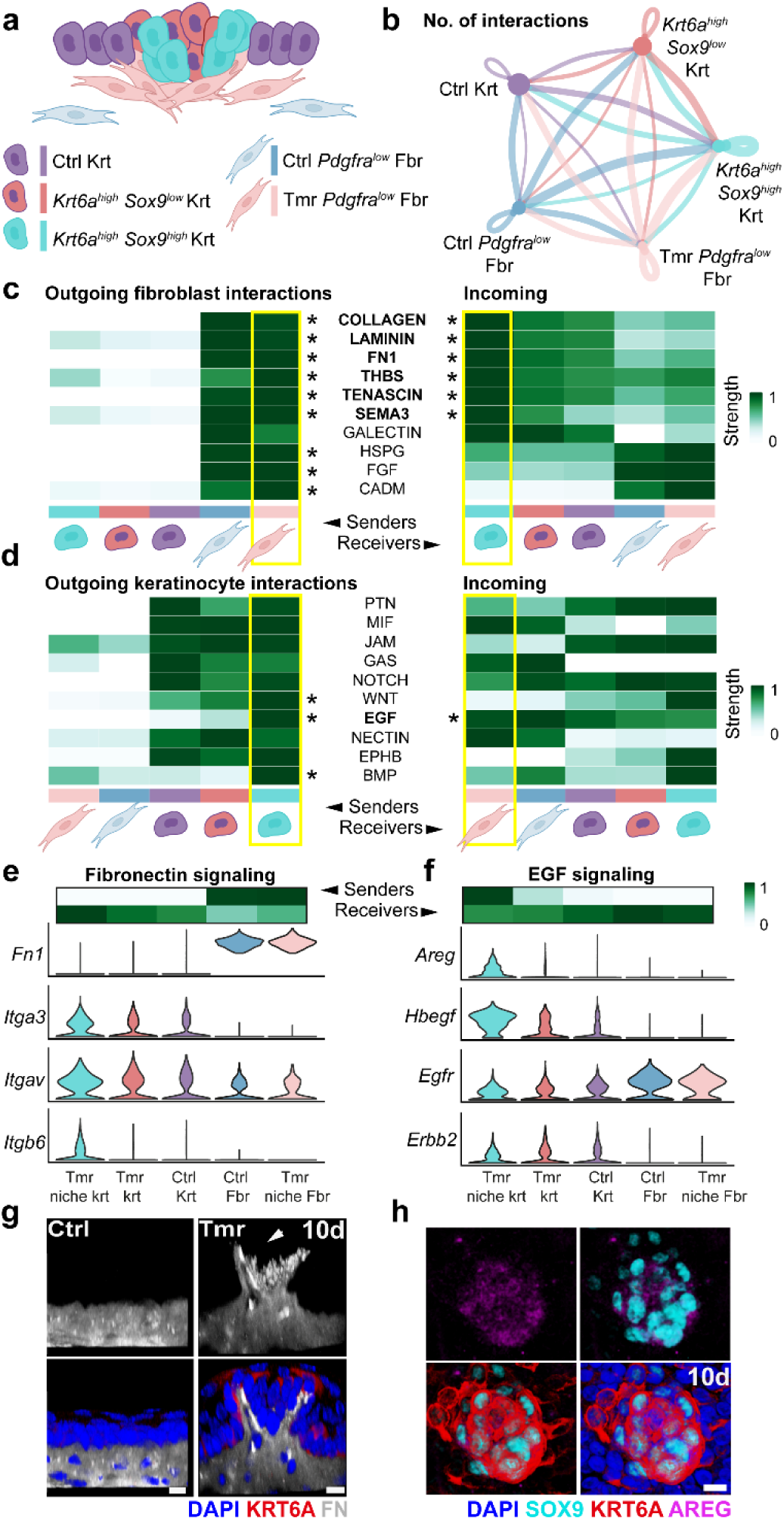
Predicted communication interactions between tumour niche keratinocytes and fibroblast reveal a EGF-Fibronectin signalling axis. **a**, Cartoon showing the heterogeneous composition of pre-neoplastic tumours. Ctrl, control; Tmr, tumour; Krt, keratinocyte; Fbr, fibroblast. **b**, Aggregated cell-cell communication network. Circles are color-coded to represent communicating cells, while circle size relates cell number; the thickness of the connections represents the number of significant interactions identified by CellChat. **c**, Heatmap showing significant top 10 outgoing fibroblast interactions (left) and matched incoming interactions (right). Tumour niche keratinocytes and fibroblasts are marked in yellow to highlight their top predicted interactions. Signals with stronger interactions predicted in tumour niche cells are marked with an asterisk. Scale is relative signal strength representing total interaction strength divided by the highest value among all cell clusters. **d,** Heatmaps equivalent to **c**, marking the significant top 10 outgoing keratinocyte interactions and matched incoming interactions. **e-f,** Heatmaps (top) showing the relative importance of each cell group based on the computed network centrality scores for the Fibronectin and EGF signalling network, respectively. Violin plots of relevant ligands and receptors across different cell types (bottom). These include expression of fibronectin (*Fn1*) and fibronectin-binding receptors (*Itga3, Itgav, Itgb6*) (**e**), as well as EGF ligands (*Areg, Hbegf*) and binding receptors (*Egfr, Erbb2*). **g**, Representative confocal 3D-rendered side-view images of a control area and nascent tumours 10 days post-DEN showing the accumulation of Fibronectin in the early tumour niche. Samples were labelled for fibronectin (FN, greyscale), KRT6A (red) and DAPI (blue). Scale bars, 10 µm. **h,** Top-down view of nascent tumour epithelium 10 days post-DEN confirming the overexpression of AREG in epithelial cells from early tumour stages. DAPI, blue; AREG, magenta; KRT6A, red; SOX9, greyscale. Scale bars, 10 µm. Panel (**a**) created with BioRender.com.

Pro-fibrotic ECM-related pathways were among the top outgoing interactions predicted to preferentially signal from tumour niche fibroblasts to tumour niche keratinocytes. These pathways included COLLAGEN, LAMININ, FN1, THBS, TENASCIN, and SEMA3 (**Fig. 5c**). The pro-fibrotic nature of tumour niche fibroblasts was further evidenced by the active ECM remodelling observed in the early tumour niche, as revealed by second harmonic generation imaging (**Extended data Fig. 9c-e**). Given its well-established role in ECM assembly and its strong association with fibrotic diseases^52,53^, we reasoned that fibronectin (FN) may represent a central player modulating ECM interactions across tumour cell compartments. The relevance of FN interactions was supported both by the specific upregulation of FN receptor genes in tumour niche keratinocytes (**receiver cells**), and by the increased expression of FN, at both mRNA and protein levels, in tumour niche fibroblasts (**sender cell**; **Fig. 5e, Extended Data Fig. 9f**). Overall, our scRNA-seq and communication analyses uncovered the pro-fibrotic/wound-healing transcriptional profile characteristic of tumour niche fibroblasts (C23). Next, we explored the outgoing signals from tumour niche keratinocytes. Here, EGF was identified as one of the strongest incoming signals for tumour niche fibroblasts (**Fig. 5d**). Indeed, the expression of EGF ligands (including *Areg* and *Hbegf*) was enriched in tumour niche keratinocytes, at both the mRNA and protein levels (**sender cell**; **Fig. 5f; Extended data Fig. 9g,h**), with its receptor (*Egfr*) being markedly expressed in the underlying tumour niche fibroblast population (**receiver cell**; **Fig. 5f**).

The upregulation of both AREG and FN in the niche of nascent tumours, observed as early as 10 days post-DEN (**Fig. 5g,h**), highlighted the significance of the EGF-Fibronectin mediated epithelial-mesenchymal cross-talk from the earliest stages of tumorigenesis.

Altogether, these data reveal the existence of an epithelial-mesenchymal EGF-Fibronectin communication axis that becomes established in the pre-cancerous tumour niche from early pre- neoplastic stages.

### EGF-Fibronectin tumour-niche communication axis mediates early tumour survival

Given the well-known role of EGF and FN signalling in epithelial-stromal communication during wound healing and tissue damage^54-56^, we hypothesized that these may be playing a similar role in response to genetic stress, such as that induced by DEN, effectively promoting tumour stroma reorganization, niche formation, tumour growth and ultimately survival.

So far, our results have shown that *Sox9*^high^ tumours are associated with stromal remodelling and niche formation (**Fig. 3**). To determine whether *Sox9*^high^ keratinocytes are exerting their mesenchymal remodelling effect via the EGF pathway, as hinted by our cell communication analysis (**Fig. 5d,f,h**), we performed a chemoattractant assay (**Fig. 6a**). We found that the EGF ligand, AREG, positively stimulated fibroblast migration, indicating that incipient tumour keratinocytes promote fibroblast chemotaxis through paracrine EGF stimulation **(Fig. 6b,c).**

**Fig. 6:**
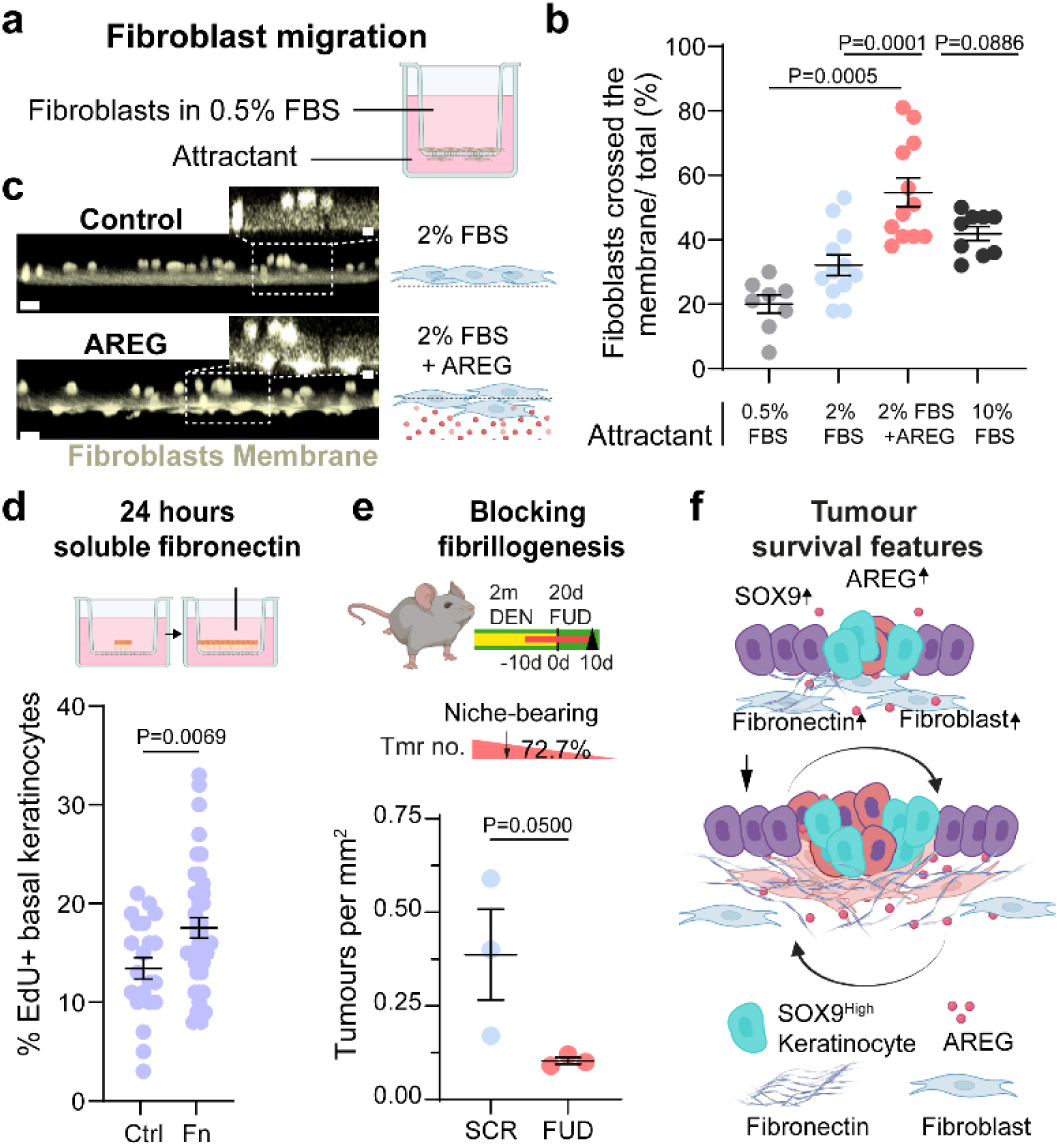
The EGF-Fibronectin axis promotes epithelial-mesenchymal communication driving fibroblast migration, keratinocyte proliferation and increased tumour burden. **a**, Fibroblasts from normal healthy tissue where plated on inserts (8.0 µm pore diameter) and exposed to different attractants in the lower insert compartment, including AREG ligand. **b**, Migration of fibroblast through the insert membrane was quantified and expressed as a percentage of the total number of cells. Data expressed as mean±s.e.m. Each dot represents a technical replicate; data 3-4 biological replicates. Brown-Forsythe and Welch’s ANOVA tests comparing AREG-supplemented condition against all others. FBS, fetal bovine serum. **c**, Representative confocal images (left), and schematic representation (right) of fibroblast migration observations. Scale bars 25 µm, 10 µm (inset). **d**, Schematic representation (top) and quantification (bottom) of EdU incorporation (2- hour chase) in confluent 3D epithelial cultures (Epithelioids) exposed to Fibronectin for 24 hours. Data expressed as a percentage of the total number of cells (mean±s.e.m). Each dot represents a technical replicate; data from 3 biological replicates. Two-tailed Welch’s t-test comparing EdU incorporation in control (Ctrl) and fibronectin-treated cells. e*, In vivo* protocol (top) and quantification (bottom) of niche+ tumours per mm^2^ in animals treated with a fibronectin assembly inhibiting peptide (FUD) for 20 days. Tumour burden decreased ∼72.7% in FUD group compared to a scrambled peptide (SCR) used as control. Tissues were collected 10 days post-DEN. n = 3 biological replicates. Mean ± s.e.m whiskers shown. One-tailed Mann-Whitney test of SCR *versus* FUD. **f**, Schematic of suggested model whereby epithelial cells respond to genetic perturbation activating a stress state marked by SOX9 and EFG ligand overexpression. This drives underlying fibroblasts to migrate towards the nascent tumour. There they remodel the pre-cancerous stroma, forming a fibrotic ECM scaffold. The established EGF-Fibronectin axis between epithelial and mesenchymal cells results in the formation of an early tumour niche that favours tumour growth and promotes long-term survival. Panels (**a**), (**b**), (**e**) and (**f**) created with BioRender.com.

Similarly, FN, a major component of the fibrotic niche in early tumours (**Fig. 5c,e,g**), represents a critical and well-established regulator of the stromal wound response^57^. FN is a structural and functional constituent of the wound granulation tissue that serves as a scaffold to support epithelial cell proliferation during tissue repair. Hence, we reasoned that the newly formed fibronectin-rich tumour niche may also promote the proliferation of tumour keratinocytes and thereby favour tumour survival. Indeed, established 3D epithelial cultures (Epithelioids)^58^ treated with soluble FN for 24 hours showed a significant increase in epithelial proliferation compared to untreated cultures (**Fig. 6d**). Accordingly, *in vivo* experiments, where animals were treated with a peptide inhibiting fibronectin fibrillogenesis (FUD)^59,60^, which interferes with matrix assembly, showed a significant reduction of niche+ tumours, hindering early tumour survival (**Fig. 6e**). We concluded that the FN signalling axis plays a critical role in promoting early tumour growth and survival by mediating the communication between the niche and the epithelium.

Our results demonstrate the central role of the EGF-Fibronectin axis in early tumour niche formation. In response to genetic stress, SOX9^high^ epithelial cells stimulate fibroblast migration through EGF signalling. This promotes the formation of a pro-fibrotic tumour scaffold, rich in fibronectin, that promotes tumour growth and survival.

## Discussion

Recent work has shown that cancer mutations, traditionally thought to be the sole cause of cancer, can also be found in healthy ageing tissues^1-5^, and form part of normal tissue physiology. This has redirected the interest of the cancer community to fill the knowledge gap on the earliest stages of the disease. Studies have recently highlighted the importance of the microenvironment, showing that the predisposition of a mutated epithelium to develop tumoral lesions able to progress towards advanced stages depends on intricate, still largely unknown, interactions between mutant cells and their dynamic surroundings^6,7,13,14,21,22,61-64^. Yet, the question remains: how does the tumour microenvironment contribute to the early and most incipient stages of tumorigenesis?

In this study, we made use of a well-established cigarette-smoke carcinogen tumour model^6,21,22^ to study the processes underlying early tumour survival. Specifically, we investigated the intrinsic differences between nascent and long-term persisting squamous tumours in the upper GI tract. Full-thickness 3D tissue reconstruction and single-cell transcriptional profiling revealed the heterogeneous nature of the nascent tumour stroma, with approximately 1 in 4 tumours presenting a marked stromal remodelling in the form of a mesenchymal structure that partially encapsulates the growing tumour. Over time, long-term persisting tumours converged towards a unique phenotype, with tumour survival being directly linked to the formation of a highly fibrotic stromal structure that acts as a pre-cancerous niche promoting tumour growth.

Transcriptional analysis within the epithelial tumour compartment revealed the presence of a *Sox9*^high^ specific tumour cell state able to promote fibroblast mobilization through EGF signalling. Functional analysis showed that the presence of the niche and the associated tumour persistence were both indeed linked to SOX9 over-expression in the epithelial tumour compartment.

We found that the reciprocal communication between mesenchymal cells residing within the tumour niche and SOX9^high^ tumour epithelial cells perpetuated tumour survival. From the earliest stages of tumour formation, niche fibroblasts contribute to ECM remodelling, forming a pre- cancerous niche that sustains tumour growth. This remodelling process involves the increased production of fibronectin, an active signalling component of the ECM^52,57,65^. Our functional *in vivo* and *in vitro* experiments proved the relevance of fibronectin signalling, stimulating tumour proliferation, promoting niche maintenance and, ultimately, favouring long-term tumour survival. Accordingly, interfering with matrix assembly by blocking fibronectin fibrillogenesis *in vivo* significantly impaired niche formation, prevented tumour survival, and reduced the overall tumour burden. Remarkably, 3D organ cultures and *in vivo* grafting experiments, in which healthy epithelium was exposed to tumour stromal cues, revealed the ability of the tumour niche, and specifically of fibronectin, to confer tumour features to a tissue that was never exposed to carcinogens, including increased epithelial cell proliferation and survival.

Our observations support a model where the fate and survival prospects of early pre-neoplastic lesions are determined by the cross-talk across cellular compartments **(Fig. 6f)**. We propose that repeated exposure to mutagens activates a “tissue stress” response whereby incipient epithelial and mesenchymal tumour cells are stimulated to signal and feedback onto each other through a SOX9-EGF-FN axis. This promotes the formation of an early tumour niche that supports tumour survival and growth. Cancer cells have recently been shown to activate CAFs via EGFR communication promoting metastasis at advanced malignant stages^66^. Similarly, EGF signalling has emerged as a key regulator of mutant clone competition at pre-cancer stages^9^. These observations argue that tumour cells may be repurposing cell communication networks to promote carcinogenesis at different stages throughout disease progression, from tumour formation to cancer dissemination. And highlights that strategies targeting both tumour cells, as well as supporting neighbouring cells may improve cancer prevention and long-term outcome.

Overall, our data reveal the unprecedented capacity of the early tumour niche to perpetuate tumour survival signals well beyond genetic alterations. This demonstrates that not only mutations but also the environmental response to genetic stress defines the likelihood of individual tumours to persist in the tissue and progress towards more advanced disease stages.

## Methods

### Mice strains

All experiments were approved by the local ethical review committees of the University of Cambridge and conducted according to Home Office project licenses PPL70/8866 and PP7037913 at the Cambridge Stem Cell Institute, University of Cambridge.

Unless otherwise specified, C57BL/6J mice (ordered from Charles River, UK; strain code 632) were used. Other mouse strains includes: cell cycle reporter line *R26^Fucci2aR^* (*Fucci2a*)^67^kindly provided by Ian J. Jackson; *Pdgfra^EGFP^* mice (stock #007669, Jackson Laboratory); *Sox9^flox/flox^* mice^68^ (obtained from MRC-Harwell, on behalf of the European Mouse Mutant Archive (https://www.infrafrontier.eu/)); *K14-Cre^ER^* mice (stock #005107, Jackson laboratory); *R26^mT-mG^* mice (*mTmG*; stock #007676, Jackson Laboratory); *Col1a2^CreER^* mice (strain #029567, Jackson Laboratory); *R26^FlConfetti^* mice^69^ (strain #017492, Jackson laboratory, kindly provided by Hans Clevers); *R26^nT-nG^* mice (*nTnG*, strain #023537, Jackson Laboratory); *H2B-EGFP* mice (*CAG::H2B-EGFP*; strain #006069, Jackson Laboratory); NOD scid gamma mice (NSG; NOD.Cg-*Prkdc^scid^ Il2rg^tm1Wjl^*/SzJ; strain #005557, Jackson Laboratory). Additional information about experimental mouse lines can be found in Supplementary Methods section ‘**Experimental mouse lines**’.

Recombination of *Col1a2^CreER^R26^FlConfetti/WT^*mice was induced by a single intraperitoneal (i.p.) tamoxifen injection (3 mg per 20 g of body weight in sunflower seed oil). *K14^CreERT^/ Sox9 ^flox/flox^* received two subcutaneous tamoxifen injections (5 mg per 20 g of body weight in sunflower seed oil) 48 hours apart.

All strains were maintained in a C57BL/6 background. All experiments comprised a mixture of male and female mice with no gender-specific differences observed (unless specified otherwise). For RNA sequencing experiments, only male animals were used in order to avoid cofounding effects due to estrous cycle. All animals exposed to the carcinogen and their respective controls were adults between 8 and 14 weeks of age (see chemically induced mutagenesis below). Mice were bred and maintained under specific-pathogen-free conditions at the Gurdon Institute and The Anne McLaren Building, University of Cambridge. All animals were housed between 20-24°C, 45-65 % humidity and a 12-hour light-dark cycle.

### Chemical tumorigenesis model

Mice were treated with Di-ethyl-nitrosamine (DEN, Sigma-Aldrich; N0756) at 40 mg per litre in Ribena- flavoured water for 24-hours, 3 days a week (Monday, Wednesday and Friday) for eight weeks^6,21^. Mice received sweetened water between DEN dosages, and normal water upon DEN treatment completion. Control mice received sweetened water as vehicle control for the length of the treatment.

### EdU lineage tracing

For EdU (5-ethynyl-2′-deoxyuridine) labelling experiments, animals received 100 µg of EdU in PBS (Life Technologies; A10044) intraperitoneally 2 hours prior to tissue collection. *In vitro* 3D cultures (see above) received media supplemented with 10 µM of EdU and incubated for 2-hours at 37 °C and 5 % CO_2_ before fixation. EdU incorporation in tissue whole-mount (see above) or was detected using Click-iT EdU kit according to the manufacturer’s instructions (Invitrogen, C10337). EdU^+^ cells were quantified using confocal microscopy.

### Fibronectin fibril assembly inhibitor treatment *in vivo*

Pharmacological inhibition of Fibronectin (FN) fibrillogenesis was achieved by treating mice with the Functional Upstream Domain (FUD) peptide^59^ intraperitoneally for 20 days at a concentration of 12.5 mg/kg body weight. Control animals were treated with scrambled (SCR) control peptide. Treatment started 10 days prior the end of DEN treatment and ended 10 days after. Peptides were synthesised at >95% purity (WatsonBio; Peptide sequence below). Lyophilised peptides were reconstituted in PBS.

FUD - Cys-GSKDQSPLAGESGETEYITEVYGNQQNPVDIDKKLPNETGFSGNMVETEDTKLN

SCR - Cys-QGQTGPVNSKVKIDNYELESNPEKIEANDLQVEGTTTYESKFMGDLTGSGNPED

### Whole-mount preparation

The upper gastrointestinal tract (oesophagus and forestomach) from control and DEN-treated mice was dissected at the indicated time points. Oesophagi were excised and cut open longitudinally. The muscle layer was then removed and tissue was flattened under a dissecting microscope using fine forceps. Stomachs were cut open longitudinally and rinsed twice with PBS to remove any food remains. The glandular stomach was excised away and the forestomach kept and flattened for downstream analysis. For whole tissue whole- mounts, the oesophagus and forestomach were fixed in 4% paraformaldehyde (PFA) (Alfa Aesar; 043368) in PBS for 30 minutes at room temperature (RT). For epithelial-only and stromal-only whole-mounts, tissues were incubated in 5 mM Ethylene-diamine-tetraacetic acid (EDTA) (Life Technologies; 15575020) in PBS for 3 h at 37°C. Following incubation, the epithelium was gently peeled from the stroma using fine forceps. Subsequently, each of these layers was individually flattened, fixed in 4 % PFA in PBS for 30 minutes at RT and stored for downstream confocal analysis.

### Histology

For histology, tissues were fixed in 10 % formalin in PBS overnight at RT prior to storage at 4°C in absolute ethanol. Hematoxylin and Eosin (H&E) staining was performed in 7 μm paraffin-embedded sections by the Histology Core Service at Cambridge Stem Cell Institute and imaged using Zeiss AxioScan Z1 microscope.

### *Ex vivo* tissue recombination assay

Under a dissecting microscope in a laminar flow hood, oesophagi were dissected and epithelial-stromal layers isolated as described in section ‘**Whole-mount preparation**’. Thereafter, tissues were rinsed in 1 % P/S in PBS three times to remove residual EDTA and flattened. Combination of epithelium and stroma from different experimental conditions (DEN treated and/or control) were prepared by carefully placing the epithelial layer over the relevant stroma (referred to as “tissue recombination” composites). The remaining epithelium was trimmed to match the size of the stroma, and the resulting construct was cut in half. Flattened epithelium-stromal constructs were cultured in 6-well plate inserts (ThinCert Greiner Bio-One; 657641). Size-matched Polydimethylsiloxane (PDMS) stencil frames were placed around the tissue construct to prevent cell expansion (see “Stencil production”). The tissue was settled for 10 min prior to adding 2 ml of minimal medium (mFAD), containing one-part DMEM (Fisher Scientific; 41966029) and one-part DMEM/F12 (Fisher Scientific; 11320033) supplemented with 5 μg/ml insulin (Sigma-Aldrich; 15500), 5 % foetal calf serum (Fisher Scientific; 26140079), 1 % P/S and 5 µg/ml Apo-Transferrin (Sigma-Aldrich; T2036), as per ^23,58^. 3D heterotypic cultures were maintained in standard humidified cell culture incubators at 37°C with 5 % CO_2_ for up to 7 days. At the end point samples were fixed in 4 % PFA in PBS for 30 minutes at RT, and stored for downstream confocal analysis.

### Stencil production

Silicone elastomer (PDMS)) was mixed with a curing agent (Avantor VWR; Sylgard 184 Elastomer Kit 634165S) at a 10:1 ratio and centrifuged at 300×g for 10 min to remove bubbles. The resulting mix was poured on a dish at around 70 mg/cm^2^ and left on an even surface to polymerise overnight at 37°C. The following day the resulting polymer was cut into 2x5 mm rectangle-shaped frames, sterilised in 70 % ethanol overnight, and treated with 1 % pluronic acid (Sigma-Aldrich; P2443-250g) in PBS for one hour at 37°C. The frames were then left to air dry prior to use.

### *In vivo* tissue recombination grafting

Tissue recombination composites (as described in section “***Ex vivo* tissue recombination assay**”) of DEN-treated oesophageal stroma and untreated (control) oesophageal epithelium were prepared for *in vivo* grafting adapting the strategy described above. Prior to separating the epithelium from the stroma, all visible tumours were marked with a partial incision using a punch biopsy tool (1 mm diameter, Merk; WHAWB100040). After separating the tissue layers, all stromal compartments were assessed for peeling efficiency under a fluorescence dissecting microscope (Leica M165 FC), and any remaining epithelium, identified by the dense epithelial nuclei clusters were excised from the tissue using a 1 mm biopsy punch. For heterotypic tissue constructs, 2 mm biopsies (Selles Medical; instrument BP20F) of tumour or controls stroma were excised and a 2 mm healthy untreated epithelium biopsy placed above. Composites were cultured overnight as described above and grafted in the back skin of anesthetized shaved NSG female mice (1-3 constructs per incision, 2 incisions per animal). Longitudinal incisions for grafting were approximately 5 mm in length. The wounds were closed with GLUture glue (Fisher Scientific; NC0632797) and the mice were left to recover. 3-6 months later, mice were culled and the back skin fixed with 4% PFA in PBS for 30 minutes at RT, and stored for downstream confocal analysis.

### Primary mouse fibroblast isolation and migration assay

Oesophagi were dissected as described above and cut in half. Tissue was the incubated in 0.5 mg/ml Dispase (Sigma-Aldrich; D4818) for 10 min at 37°C rotating. After incubation, the epithelium was peeled away, and the stroma was minced finely and incubated in Trypsin-EDTA (0.25%) (ThermoFisher; 25200056) for 15 min at 37°C rotating. The resulting suspension was mixed by pipetting and DMEM supplemented with 10 % FBS and 1 % P/S was added (1:1 vol:vol). The suspension was passed through a 70 µm filter (PluriSelect; 43-10070- 40) and cells pelleted by centrifugation at 300×g for 5 min at 4°C. Pellets were resuspended in 0.5 % FBS, 1 % P/S in DMEM, and seeded on 8.0 µm pore transwell insert (24-well plates; ThinCert Greiner Bio-One; 662638). The lower compartment of the transwell contained 1 % P/S DMEM supplemented either with 0.5 % FBS, 2 % FBS, 10 % FBS or 2 % FBS with 1 g/ml Amphiregulin (AREG) (R & D; 989-AR-100/CF). Primary fibroblasts were cultured for 48 hours prior to fixation in 4 % PFA in PBS for 10 min. Membranes were incubated with 1 µg/ml DAPI (Sigma-Aldrich; D9542) in PBS for 30 min at RT, cut and mounted in 1.52 Rapiclear mounting medium (SUNJin Lab; RC152001) keeping their original orientation, followed by confocal analysis. Further information on quantification can be found in Supplementary Methods section ‘**Analysis of fibroblast migration assay**’.

### Keratinocyte cultures and fibronectin treatment *in vitro*

Oesophagi were cultured using the 3D epithelioid organ culture approach^58^. Briefly, tissues were dissected as described above, cut into 3x5 mm rectangles, and placed on a transwell insert with the epithelium side up. Tissue was left to settle for about 5 min. Explants were expanded in complete medium (cFAD) containing mFAD supplemented with 1x10^−10^ M cholera toxin (Sigma-Aldrich; C8052), 10 ng/ml Epidermal Growth Factor (EGF, Fisher Scientific PeproTech^R^; AF-100-15), 0.5 μg/ml hydrocortisone (Calbiochem; 386698). Tissue explants were removed by aspiration five days after culture set up and maintained in mFAD for 2 weeks to confluence. Soluble fibronectin (Fisher Scientific Corning ™; 356008) was added to the medium for 24 hours at 100 μg/ml after diluting it in mFAD with 25mM Hepes (Fisher Scientific; 15630056). Samples were fixed with 4 % PFA in PBS for 30 minutes at RT, after a 2h EdU chase, and kept for downstream confocal analysis.

### Immunostaining

After fixation, epithelial-stromal composites were incubated for 30 min in permeabilisation buffer (PB1; 0.5 % bovine serum albumin (VWR International, 126575-10), 0.25 % fish-skin gelatin (Sigma-Aldrich; G7765), 1 % Triton X-100 (Fisher Scientific Ltd, 10102913) in PBS), then blocked for 2-hours in PB1 containing 10 % donkey serum (DS) (Scientific Laboratory Supplies, D9663). Next, tissues were incubated with primary antibodies diluted in 10% DS in PB1 for 3 days at 4°C followed by four washes over the following day with 0.2% Tween-20 (Promega UK Ltd, H5151) in PBS. Thereafter, tissues were incubated overnight with secondary antibodies diluted 1:500 in 10 % DS in PB1 at RT. Unbound antibody was removed by four washes with 0.2 % Tween-20 in PBS throughout the next day. Antibody details are provided in **Supplementary Table 1**. To stain cell nuclei, tissues were incubated with 1 µg/ml DAPI in PBS at 4°C overnight. Afterwards, samples were rinsed three times in PBS and mounted in 1.52 Rapiclear mounting media for imaging.

Immunolabeling of individual tissue layers (epithelium or stroma) consisted of an incubation for 30 min in permeabilisation buffer (PB2; 0.5 % Bovine serum albumin, 0.25% Fish skin gelatin, 0.5 % Triton X-100 in PBS). Tissues were then blocked for 2-hours in PB2 containing 10% DS. Next, samples were incubated overnight at RT with primary antibodies diluted in 10% DS in PB2 followed by three washes with 0.2 % Tween-20 in PBS throughout the next day. Secondary antibodies were diluted 1:500 in 10 % DS in PB2 and incubated with tissues overnight at 4°C after which unbound antibody was removed by three washes with 0.2 % Tween-20 in PBS and staining continued as above. Thick cryosections of fixed tissues embedded in optimal cutting temperature compound (OCT; Thermo Scientific; 12678646), cut with a thickness of 50 µm onto glass slides, were immunolabelled using the exact same protocol.

When staining with primary antibodies raised in the same host, one of these antibodies was acquired as pre-conjugated with a fluorophore or conjugated in-house following manufacturer’s instructions (Invitrogen; A20186/A20187). The staining proceeds as described above with the unconjugated antibodies. After incubation with the corresponding secondary antibodies, the samples were blocked for 3 hours at RT with 10 % DS in PB with the IgG from the relevant host species (1:500). Afterwards, samples were incubated with conjugated antibodies diluted in PB containing 10 % DS and the relevant host IgG (1:500) overnight at RT. At this point staining proceeds as above. Immunostained samples were analysed by confocal imaging.

### Confocal Imaging

Confocal images were acquired using either an inverted Leica SP5 microscope with standard laser configuration, or a Stellaris 8 FALCON FLIM microscope with a white light laser. Typical confocal settings used included: bidirectional scanning, a 40× immersion objective, an optimal pinhole size (as defined by the software), a scan speed of 400-600 Hz with 2-3x line averaging, optimal Z-step size (as defined by the software) and a resolution of 512 × 512 or 1,024 × 1,024 pixels unless stated otherwise. Three-dimensional reconstructions from optical sections and their corresponding image renders were generated using Volocity 5.3.3 software (PerkinElmer), Zen 3.2 and Arivis 3.5.1. Further information about specific image analysis can be found in **Supplementary Methods** sections.

### Second-harmonic generation (SHG) imaging

Extracellular matrix (ECM) fibres in whole-mounts were imaged by SHG microscopy in a Zeiss LSM880 microscope. Prior to imaging, fixed oesophagi were incubated in PBS with 10 drops of NucRed647 (nuclear staining; Life Technologies, R37106) for 1 h at RT to label cell nuclei. Immediately after, samples were mounted using RapiClear 1.52 medium. For image acquisition, a Plan-Apochromat 40×/1.3 Oil DIC M27 objective was used. Tumours were identified by their characteristic morphology using a 633 nm laser to excite the NucRed647 nuclear dye, detecting its emission with a 638-704 nm filter. After the region of interest was defined, the tuneable laser of the instrument was set to 1040 nm (gain set to 780) to generate second-harmonic light from the ECM fibres in the tissue, which was detected with a 481-526 nm filter. Nuclei and ECM fibres were each imaged independently for every region of interest and merged afterwards for visualization using the ZEN software (Blue edition, version 3.1, Carl Zeiss Microscopy GmbH). The microscope pinhole was set to its maximal aperture, namely 600 μm, and 2× line averaging was used. Images were acquired every 1 µm along the Z axis with an XY resolution of 1024 x 1024 pixels, a depth of 12 bits per pixel, and a pixel dwelling time of 0.77 µs. The information about SHG image analysis can be found in Supplementary Methods section ‘**SHG image analysis**’.

### Transcriptomics

#### Single-cell and RNA isolation for single-cell RNA-Sequencing (scRNA-Seq)

For scRNA-Seq, *Fucci2a* male mice were sacrificed 8 months after DEN treatment (age-matched vehicle treated mice were used as controls). After dissection, as described above, tissues were flattened with the epithelial side up and visible tumour lesions were marked by a partial incision using a punch biopsy tool (1 mm diameter) under the dissecting microscope. Then, oesophagi and forestomachs were cut in half and incubated in 50 mM EDTA in PBS containing 1 U/µl RNAse Inhibitor (Life Technologies; AM2696) at 37°C for 30 minutes. Following incubation, the epithelium was carefully peeled away from the stroma using fine forceps. Epithelium and stroma were each flattened, and tumour areas fully excised using 0.5-2 mm punch biopsies (to fit tumour size). Epithelium and stroma biopsies were kept separated. Biopsies were pooled from two to four animals. Stroma was thoroughly minced and incubated in 4 mg/ml collagenase type 4 (Thermo Fisher Scientific; 17104019) in DMEM with 1 U/ml RNAse inhibitor for 1 h at 37°C in a rotator. EDTA was added to the samples at a final concentration of 5 mM and the suspension was diluted 1:5 in collection buffer (2 % heat-inactivated fetal bovine serum, Life Technologies; 26140079; 25 mM HEPES, Life Technologies; 15630056) in PBS to reduce the collagenase activity. Cell suspensions were filtered through a 100 µm cell strainer, centrifuged at 300×g for 5 min at 4°C, and resuspended in collection buffer before an additional filtering step with a 40 µm FlowMi tip filter (Thermo Fisher Scientific; 15342931) was performed to obtain single-cell suspensions. Cells were centrifuged at 300×g for 5 min at 4°C and resuspended in collection buffer containing 1U/µl RNAse. In parallel, the epithelium was thoroughly minced and incubated for 10 min at 37°C in 0.5mg/ml dispase (Sigma-Aldrich; D4818) in PBS containing 1 U/µl RNAse to create an epithelial cell suspension. EDTA was then added to the samples at a final concentration of 5mM and the suspension diluted 1:5 in collection buffer to reduce dispase activity. A single-cell suspension was obtained by filtering the samples through a 30 µm cell strainer. Cells were centrifuged at 300×g for 10 min at 4°C and resuspended in collection buffer containing 1 U/µl RNAse. Epithelial and stromal cells were then pooled together by condition (Control, Ctrl; Adjacent non-tumour epithelium, DEN; and Tumour) prior to proceeding to library production for scRNA-seq.

#### Library preparation and scRNA-Seq

Single-cell RNA-seq libraries were generated using the 10x Genomics Chromium Next GEM Single Cell 3’ Reagent Kit (v3) and sequenced at the Genomics Core Facility of Cancer Research UK (CRUK), Cambridge Institute. Libraries were generated in two different batches. Control samples were included in both batches to provide a reference to assess potential batch effects. The cells for each biological replicate were loaded into a 10x Chromium microfluidics chip channel to generate one library from each. In total, 17 libraries were sequenced on either an Illumina HiSeqx4000 or a NovaSeq6000 system using one SP, two S1, and two S2 flow cells. Of note, given the punch biopsy approach used, DEN samples may contain sporadic tumour cells from tumours not visible to the naked eye.

#### scRNA-seq data pre-processing, dimensionality reduction, and visualization

The raw scRNA-Seq data was processed using CellRanger (version 3.0.2). CellRanger aligned the reads to the mouse reference genome (Ensembl GRCm38/mm10; release 92), filtered empty droplets, and counted unique molecular identifiers (UMIs) to generate gene expression matrices. To filter out low-quality cells, basic quality control metrics were calculated using the R package scater^70^ (1.12.2), and cells with gene counts below 400 or above 8950 were removed. Cells with over 15 % of reads mapping to mitochondrial genes, as well as genes expressed in fewer than three cells, were not considered for downstream analyses. Read counts were subsequently normalised via a deconvolution method using the R package scran^71^ (version 1.12.1). The normalised data was grouped by tissue of origin (i.e., oesophagus or forestomach). Principal component analysis, combined with technical noise modelling, was applied to each of the split datasets for dimensionality reduction, as implemented in the denoise_PCA function of the R package scran. The data were then projected in two dimensions by using the runUMAP function of the R package scater with default parameter settings. The biological replicates for the same condition overlapped well with each other, confirming negligible batch effects; therefore, no batch effect correction was necessary. The R package Seurat^72^ (version 3.0.2) was used to visualise cells in the dimensionally reduced space. The sequencing dataset was deposited at the Gene Expression Omnibus (GEO) under dataset accession number.

#### Data clustering and cluster annotation

Cell clusters were identified using the buildSNNGraph function of the R package scran, which deploys the Louvain community detection method^73^; oesophageal and forestomach clusters were identified separately. Cell clusters were annotated based on differentially expressed genes (DEGs) and on the expression level of known marker genes for the different cell types forming the tissues. In the case of multiple neighbouring clusters sharing key gene expression patterns, such clusters were manually assigned the same cellular identity. First, all clusters were classified into three broad cell groups based on the expression of marker genes of keratinocytes (e.g. *Krt13, Krt14*), stromal cells (e.g. *Vim, Pdgfra*), and immune cells (e.g. *Ptprc*). During the annotation process, pseudo cells (i.e. not real cells, but droplets with ambient RNA) and doublets (i.e. droplets with more than two cells) were identified based on two criteria: 1) the co-expression of marker genes from two different groups (e.g. keratinocytes and stromal cells), and 2) total number of genes (i.e. ≤1500 genes). Cells meeting either of these two criteria were excluded from further analysis. 66236 cells remained in the dataset.

To further investigate keratinocytes and stromal cells from the upper gastrointestinal tract, we decided to perform analyses on a merged dataset containing only these two major cell groups. Within the dataset, we defined three main conditions: age-matched control (Ctrl), DEN-treated but morphologically normal tissue (DEN), and Tumours from DEN treated mice. No obvious separation in distribution of Ctrl, DEN or tumour conditions was found (**Extended data Fig. 4c**). By repeating the dimensionality reduction process and clustering (as explained above) with a k-value (nearest neighbour) of 11, we obtained *38* cell clusters in the resulting two-dimensional representation. These clusters were annotated to 11 known biological cell types and one transitional cell type using canonical marker genes. A heatmap depicting the most representative cell markers was generated from the values obtained by extracting the average expression value of each gene for each of the clusters and scaling such values in a gene-wise manner to a range from -3 to 3. The natural-log-normalised expression levels of typical marker genes were visualised in UMAP space using the Seurat function FeaturePlot. The enrichment of early tumour markers (*Krt6a* and *Krt17*) in tumour samples validated our punch biopsy method as a useful approach to study the molecular traits of early tumour lesions (**Extended Data Fig. 6a,b**).

#### Cell transition trajectory analysis

Monocle 2^74^ (version 2.10.1) was used to order basal keratinocytes (Clusters 1, 13, 19, 21, 24, 28, 36; number of cells Ctrl=4138, DEN=1027, Tumour=3536; Total= 8701) along a pseudotime trajectory. Cell ordering was performed based on the top 1200 DEGs (ranked by lowest to highest Benjamini-Hochberg-corrected p-value, i.e., q-value) identified across conditions by using the differentialGeneTest function of Monocle with the fullModelFormulaStr parameter set to “∼condition1”. Default monocle parameter settings were used.

Monocle ordered the cells along an inferred pseudotime (PST) trajectory with 3 states and a single branching point. To determine how the cells placed on the trajectory from State (S) 1 to S2 (S1S2) differed from those placed on the S1S3 trajectory, we performed Branched Expression Analysis Modelling (BEAM) in Monocle using the BEAM function with the branch_point parameter set to 1. The top 1000 differentially expressed genes identified by the BEAM function were then plotted in a heatmap using the plot_genes_branched_heatmap function of Monocle with the num_clusters parameter set to 16 and the branch_point parameter set to 1 (**Extended Data Fig. 5c**). To assess the patterns of expression by which the genes were clustered in such heatmap, smooth spline curves were fitted along each the S1S2 and S1S3 trajectories for the expression level of each of the genes using the genSmoothCurves function of monocle with the trend_formula parameter set to “∼sm.ns(Pseudotime, df=3) * Branch”. Then, the obtained values were log_10_-transformed, averaged for all the genes in each cluster and trajectory (S1S2, S1S3) using the rowMeans fuction, and plotted with ggplot2 (v3.5.0) with minimum and maximum y-axis values adjusted for each of the clusters (identical for each of the two trajectories). Patterns are shown to the right of the heatmap in **Extended Data Fig. 5c.** The information about DEGs analysis can be found in Supplementary Methods section ‘**Differential gene expression and gene set enrichment analysis**’.

#### Analysis of interactions between cell types by CellChat

To quantitatively analyse cell-to-cell communication networks from scRNA-seq data we used CellChat^51^ (v1.1.3), a tool that infers ligand-receptor interactions (http://www.cellchat.org/). For the DEN and LE conditions, we first selected relevant keratinocyte clusters (13, 19, 21, 24, 28, and 36) and fibroblasts from cluster 23 (enriched in tumour samples). We then set thresholds for normalised expression of 1.0 for *Krt6a* and 0.3 for *Sox9*. This approach yielded five groups of cells, namely: 1) *Krt6a^high^Sox9^high^*tumour niche keratinocytes (n=701), 2) *Krt6a^high^Sox9^low^*tumour keratinocytes (n=1464), 3) *Krt6a^Low^* DEN control keratinocytes (n=1877), 4) tumour niche fibroblasts (n=836), and 5) DEN control fibroblasts (n=257). Using these five groups as input, the significant ligand-receptor interactions were identified and quantified based on CellChat’s function for computing communication probabilities for overexpressed ligand-receptor pairs using default parameter settings (trimean expression value for each group, p-value < 0.05). Total interaction strengths represent the sum of all incoming and outgoing communication probabilities for each cell cluster (and for each given pathway, if applicable). Scores for sender and receiver roles of cell clusters were computed from network centrality scores implemented through CellChat’s netAnalysis_computeCentrality function.

### Low input targeted DNA sequencing

Oesophagi from control and DEN-treated animals were dissected as described above. Tissues were flattened with the epithelial side up and visible tumour lesions were marked with a partial incision using a 1mm diameter punch tool under a dissecting microscope. Tissues were then incubated in 5mM EDTA for 3 hours at 37°C rotating. After incubation, the epithelium was removed, the stroma was flattened, and tissues were fixed as described above.

Immunofluorescent labelling against KRT14 was performed as described above. Only tumour stroma footprints negative for KRT14 (i.e. lacking epithelial cells) were considered for DNA sequencing in order to avoid the identification of genetic mutations present in epithelial cells. Tumour stroma matching the criteria was dissected under a fluorescence dissecting microscope (Leica M165 FC) using a 1 mm punch biopsy tool. Punch biopsies of equivalent size were collected from untreated healthy tissues as control. The DNA from individual biopsies was extracted using the Arcturus®PicoPure® DNA Extraction Kit (Fisher Scientific Ltd, KIT0103) following the manufacturer’s instructions. Briefly, Proteinase K was reconstituted in 155 µl and the sample was lysed in 20 µl at 65°C overnight. Thereafter, Proteinase K was inactivated by incubation at 95°C for 10 min.

Samples were then sheared, libraries prepped using the NEBNext®Ultra™ II Fragmentase System, and index tags applied (Sanger 168 tag set). Material was subjected to 12 PCR cycles (Initial denaturation; 95°C, 5 min; 12 cycles: 98°C, 30 sec; 65°C, 30 sec; 72°C, 2 min; final elongation: 72°C, 10 min) and quantified (Accuclear dsDNA Quantitation Solution, Biotium). 500 ng of pooled material was taken forward for hybridization capture and enrichment (SureSelect Target enrichment system, Agilent technologies) using a previously designed bait panel of 192 genes, including those commonly mutated in squamous cancers^22^. After cleanup, libraries were normalised to ∼6nM and submitted to cluster formation for sequencing on a Novaseq6000 (Illumina) to generate 100 bp paired-end reads.

Aligned reads were mapped to the mouse GRCm38 reference genome using BWA-mem (version 0.7.17)^75^. Duplicate reads were marked using SAMtools^76^ (v1.11). Depth of coverage was also calculated using SAMtools to exclude reads which were: unmapped, not in the primary alignment, failing platform/vendor quality checks, or PCR/Optical duplicates. Bedtools (version 2.23.0)^77^ was then used to calculate the depth of coverage per base across samples.

Variant calling was performed using the deepSNV R package (also commonly referred to as ShearwaterML; version 1.21.3; https://github.com/gerstung-lab/deepSNV). Variants were annotated using VAGrENT. R version 3.3.0 was used (from 2016-05-03)^4,78^.

Mutations called by deepSNV ShearwaterML were filtered using the following criteria: 1) positions of called SNVs have a coverage of at least 100 reads, 2) germline variants called from the same individual are omitted from the list of called variants, 3) adjustment for FDR and mutations use support from at least one read from both strands for the mutations identified, 4) pairs of SNVs on adjacent nucleotides within the same sample are merged into a dinucleotide variant if at least 90% of the mapped DNA reads containing at least one of the SNV pairs also contained the other one. DeepSNV ShearwaterML was run with a normal panel of approximately 12k reads. The sequencing dataset was deposited at the European nucleotide Archive (ENA) under dataset accession number ERP134942.

### Statistics and reproducibility

The numbers of biological replicates and animals are indicated in figure legends (n refers to the number of independent replicates per time point and/or condition). A minimum of three independent mice or *ex vivo* cultures were used in all cases. Results were consistent across replicates. For image analysis, a minimum of three independent samples were inspected in all cases. All figures show representative images. The data are expressed as mean values ± s.e.m. unless otherwise indicated.

Differences in tumour burden were assessed by one-tailed unpaired non-parametric Mann-Whitney U tests. For large datasets normality was assessed by Kolmogorov Smirnov test; for normally distributed data, differences between two groups were assessed by two-tailed Welch’s t-tests; for not normally distributed data two-tailed Mann Whitney U test was used. Differences between more than two groups were calculated using either one-way Welch’s ANOVA, followed by a Dunnett’s T3 multiple comparison test or Kruskal-Wallis one-way ANOVA, followed by Dunn’s multiple comparison test, for normally distributed or not normally distributed data, respectively. Exact P-values are indicated in the relevant figures with a precision of up to four decimal places. Statistical tests were conducted in GraphPad Prism (10.1.2) with 95% confidence interval. No statistical method was used to predetermine sample size. The experiments were performed without randomization. Blinding was performed for tumour count per condition and *in vitro* sample analyses by confocal microscopy. In cases were quantification was performed in tumours and the morphologically normal areas blinding was not possible due to differences in physical sample appearance.

## Acknowledgements

We thank members of the Alcolea’s lab for comments and suggestions; Peter Humphreys and Darran Clements in imaging core facilities at Jeffrey Cheah Biomedical Centre (JCBC); Nicola Lawrence at the Gurdon Institute Imaging Facility; Irina Pshenichaya in Histology facility at JCBC for technical support; Betania Mahler-Araujo and James Warner from Histopathology at Institute of Metabolic Science-Metabolic Research Laboratories Medical Research Council Metabolic Diseases Unit for histopathology analysis; Maike Paramor and Vicky Murray in Next Generation Sequencing Core Facility at JCBC; Katarzyna Kania in Single Cell Team at Cancer Research UK Cambridge Centre Cancer Institute; Irina Mohorianu’s team at the bioinformatics facility at JCBC for her contribution to the scRNA-seq support; the staff of the University Biomedical Services Gurdon Institute and The Anne McLaren Building, as well as the other JCBC core facilities. We are also grateful to Prof. Ian J. Jackson (Fucci2a) and Prof. Hans Clevers (Rosa26Confetti) for donation of mouse lines. Parts of the figures were Created with BioRender.com. This work was mainly supported by grants to M.P.A from the Wellcome Trust and The Royal Society (105942/Z/14/Z and 105942/Z/14/A), Medical Research Council (MR/P019013/1), Worldwide Cancer Research (19-0192 and 23-0063). This research was funded in whole, or in part, by the Wellcome Trust [203151/Z/16/Z, 203151/A/16/Z] and the UKRI Medical Research Council [MC_PC_17230]. For the purpose of open access, the author has applied a CC BY public copyright licence to any Author Accepted Manuscript version arising from this submission. G.S was funded by the Isaac Newton Trust (21.07(a)), Medical Research Council (MR/P019013/1), Worldwide Cancer Research and Guts UK (19-0192 and 23-0063). J.E.R.A was supported by a University of Cambridge / Wellcome Trust Junior Interdisciplinary Fellowship (ISSF 11/2/2020), and the Medical Research Council (MR/P019013/1). Y.D. was funded by an ELBE Postdoctoral Fellowship from the Center for Systems Biology Dresden (CSBD). M.T.B received funding from the European Union’s Horizon 2020 research and innovation programme under the Marie Sklodowska-Curie grant agreement No 794664 (OESOPHAGEAL FATE). M.T.B was also supported by the Isaac Newton Trust (Research Grants 16.24(e)) and Leverhulme Trust (RPG-2023- 136). S.H. acknowledges funding from the Human Frontier Science Program (LT000092/2016-L). PHJ and JCF are supported by the Wellcome Trust (grant 108413/A/15/D) and a Cancer Research UK programme grant (C609/A27326). This project has received funding from the European Research Council (ERC) under the European Union’s Horizon 2020 research and innovation program (grant agreement no. 950349) awarded to S.R. B.D.S. acknowledges funding from the Royal Society (E.P. Abraham Research Professorship, RSRP\R\231004) and Wellcome (098357/Z/12/Z and 219478/Z/19/Z).

## Author contributions

G.S., J.E.R.A. and M.P.A. designed, validated and conducted experiments; S.H. guided the experimental design for single-cell RNA sequencing and together with J.E.R.A. performed data processing. J.E.R.A. conducted the trajectory inference analysis. Y.D. carried out cell-to-cell communication analysis and overall advised and supported single-cell RNA sequencing analysis. S.R. analysed lineage tracing data and provided expert advice and guidance on single-cell RNA sequencing data analysis. J.C.F. processed DNA sequencing samples and performed data analysis. B.C. provided insights and technical expertise in targeted DNA sequencing. M.T.B. carried out *in vivo* transplantation assays. M.T.B., B.C., P.H.J. and B.D.S advised on various parts of the study and provided expertise in the epithelial stem cell and tumour biology fields and assisted with manuscript writing. G.S. and M.P.A conceived the project, supervised/performed experiments, and wrote the manuscript with input from all authors. Review & Editing of final manuscript by all authors.

