## supplementary figures and methods for "Pre-cancerous Niche Remodelling Dictates Nascent Tumour Survival"

#### Extended Data Figures and Legends

**Extended Data Fig. 1:** Related to Fig. 1

**Extended Data Fig. 2:** Related to Fig. 1

**Extended Data Fig. 3:** Related to Fig. 2

**Extended Data Fig. 4:** Related to Fig. 3

**Extended Data Fig. 5:** Related to Fig. 3

**Extended Data Fig. 6:** Related to Fig. 3

**Extended Data Fig. 7:** Related to Fig. 4

**Extended Data Fig. 8:** Related to Fig. 4

**Extended Data Fig. 9:** Related to Fig. 5

#### Supplementary Information

**Supplementary Table 1.** Antibodies list

#### Extended Data Figure 1

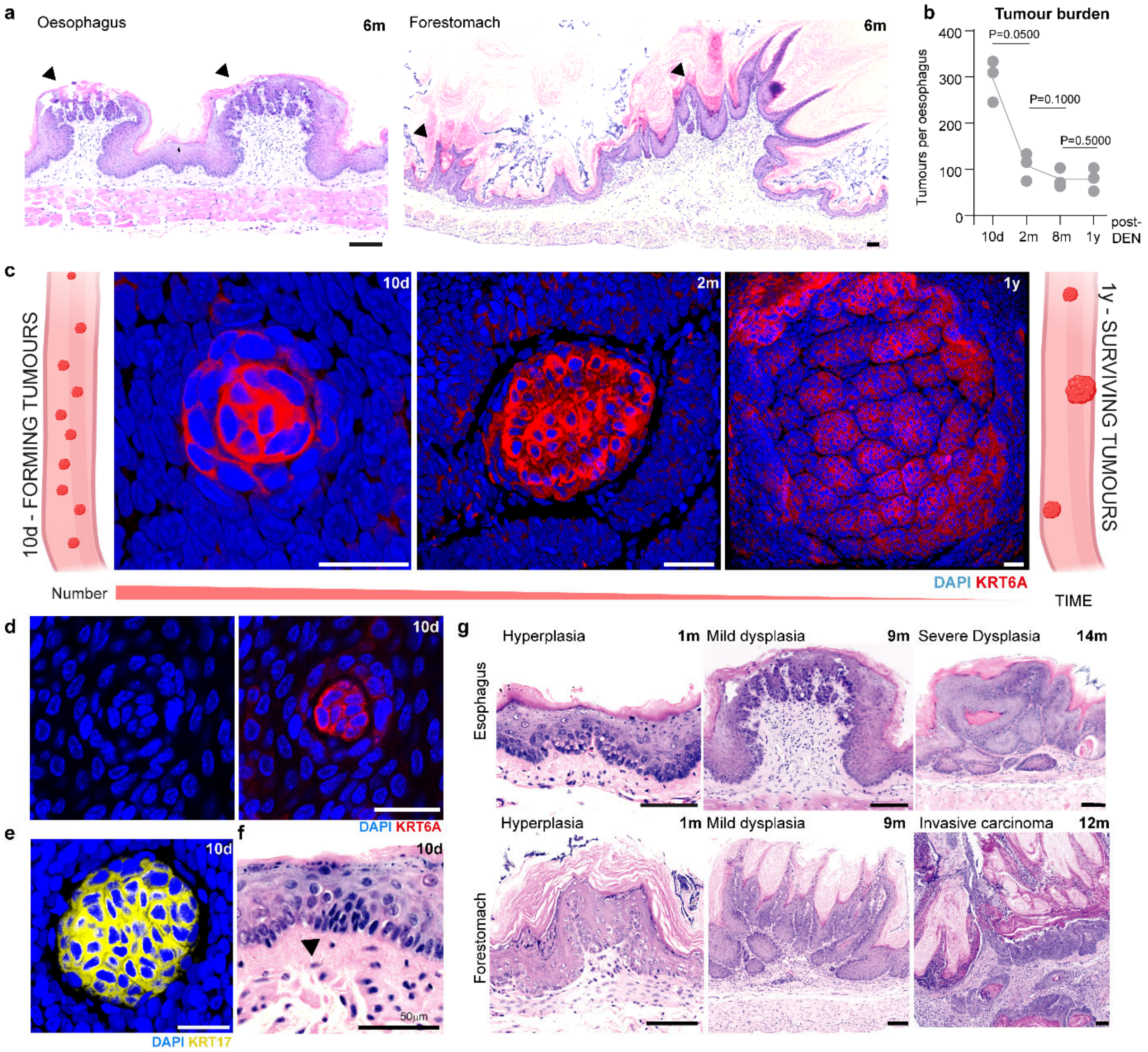


**Extended Data Fig. 1: Early tumourigenesis model in squamous gastrointestinal tract of a mouse; (Corresponds to Fig. 1). a,** H&E image of a transversal cross-section from the upper gastro-intestinal tract (including oesophagus and forestomach) displaying multiple tumours (black arrowheads) 6 moths (m) post-DEN. Scale bars, 100 µm. **b**, Number of tumours per oesophagus (OE) at the indicated time points post-DEN treatment, n = 3 mice per time point. A line is drawn across the means of each time point. For pairwise comparisons, a one-tailed Mann-Whitney test was used. **c**, Cartoon (edges) illustrating tissue tumour clearance over time, with examples of confocal images (middle) of growing persistent/surviving tumours stained for DAPI (blue) and KRT6A (red). Tissues were collected at the indicated time points post-DEN from 10 days (d) to 1 year (m). Scale bars, 25 µm. **d-e,** Confocal images showing early tumour marker expression in nascent tumours 10 days post-DEN. DAPI, blue; KRT6A, red (**d**) and KRT17, yellow (**e**). Scale bars: 25 µm. **f**, H&E image of transversal cross-section depicting a nascent tumour 10 days post-DEN, indicated by black arrowhead. **g**, H&E images of transversal cross-sections from tumours at different stages of progression in the squamous upper gastro-intestinal tract at different time points post-DEN. Scale bars, 50 µm. Panels (**c**) created with BioRender.com.

#### Extended Data Figure 2

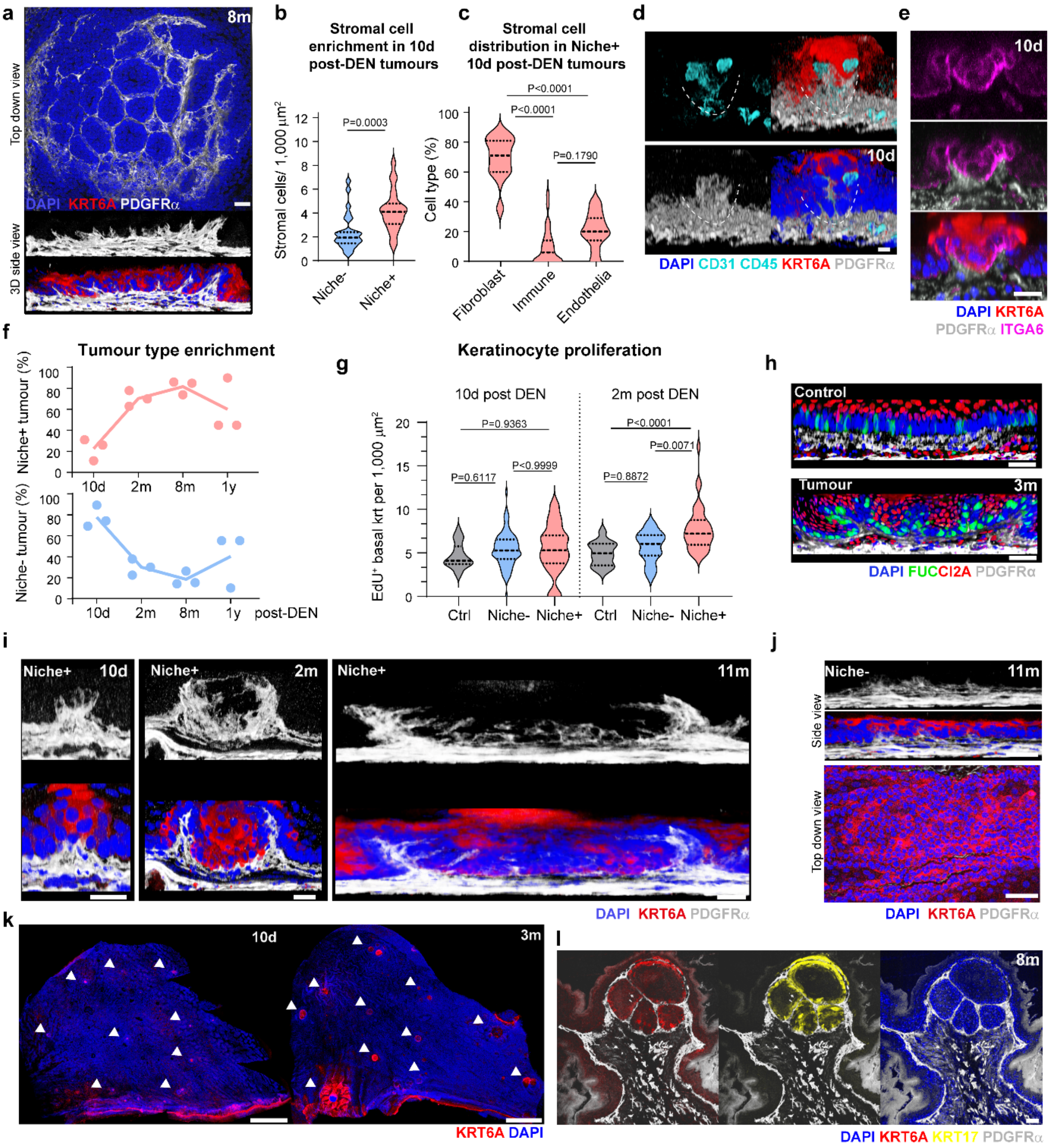

**Extended Data Fig. 2: Early tumour niche is mainly comprised of fibroblast; (Corresponds to Fig. 1). a**, Representative confocal image of a long-term persisting/surviving oesophageal tumour 8 months (m) post-DEN. DAPI, blue; KRT6A, red; PDGFR$\alpha$, greyscale. Scale bar, 25 µm. **b**, Number of stromal cells in niche- and niche+ tumours per surface area, 10 days post-DEN. n=27 (niche-), 22 (niche+) tumours, respectively, from 3 mice. Two-tailed Mann Whitney was used to asses significance. **c**, Stromal cell subtypes found in the nascent tumour niche expressed as a percentage of total stromal cells. Kruskal-Wallis ANOVA tests were used to assess differences in cell type fractions. n = 22 tumours from 3 animals. **d**, Representative confocal images showing the stromal cell composition of the nascent tumour niche 10 days (d) post-DEN. CD45+ immune cells and CD31+ endothelia cells, cyan; KRT6A tumour cells, red; PDGFR$\alpha$+ fibroblasts, greyscale. Distribution and enrichment in KRT6A-labelled niche+ tumours. Dashed white line labels tumour stromal niche. Scale bar, 10 µm. **e**, Side-view of Z-stack of images of niche+ tumour labelled for ITGA6, PDGFR$\alpha$, KRT6A and DAPI (scale bar, 10 µm). **f**, Number of niche+ and niche- tumours expressed as a percentage of total tumours at indicated time points post-DEN (m, month; y, year). n = 3 mice per time point. A line is drawn across the means of each time point. **g,** Number of EdU+ basal keratinocytes per tumour normalised per area at the indicated time points post-DEN. n- number of normal areas or niche- or niche+ tumours; n=84 and 20 (niche- tumours), n=35 and 38 (niche+ tumours), and n=20 and 14 (equivalent control areas) for 10 days and 2 months post-DEN, respectively. 3 mice per condition. Brown-Forsythe and Welch's ANOVA tests were used for multiple comparison. **h**, 3D-rendered confocal side-views of a tumour 3 months (m) post-DEN and its respective age-matched control showing $R26^{Fucci2aR}$ tissue (mCherry, G1 cells; red mVenus, S/G2/M cells; green). PDGFRα labels fibroblasts in greyscale. Corresponds to **Fig. 1f**. Scale bar, 50 µm. **i**, 3D-rendered confocal side-views showing growing niche+ tumours at the indicated time points post-DEN; DAPI, blue; KRT6A, red; PDGFR$\alpha$, greyscale. **j**, Equivalent image to **i** showing niche- tumour 11 months post-DEN, for comparison. Scale bars, 25 µm (**i** and **h**). **k**, Representative confocal images of forestomach epithelia 10 days and 3 months post-DEN. DAPI, blue; KRT6A, red. White arrowheads highlight tumours at indicated time points. Scale bars, 1.5mm. **l**, Representative confocal images of a transversal cross-section of a forestomach tumour labelled showing the presence of the stromal niche 8 months post-DEN. DAPI, blue; KRT6A, red; KRT17, yellow; PDGFR$\alpha$, greyscale. Scale bar, 25 µm. Data in panels **b**, **c** and **g** shown as violin plots with mean (solid lines) and quartiles (dashed lines).

#### Extended Data Figure 3

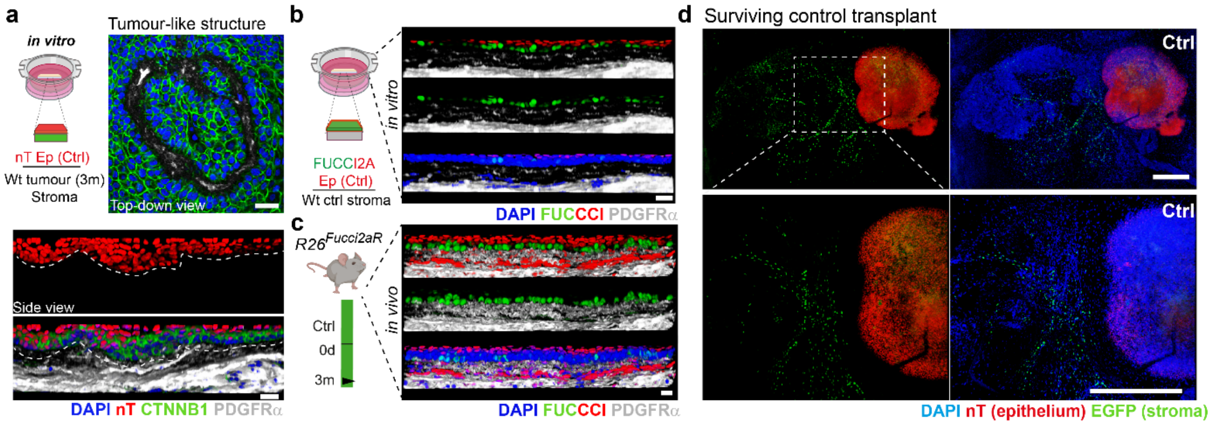


**Extended data Fig. 3: The tumour stromal niche promotes keratinocyte survival; (Corresponds to Fig. 2) a**, A schematic representation of the heterotypic construct approach used (top left). After peeling off epithelial and stromal layers, healthy untreated epithelium was combined with stroma of surviving tumours 3 months (m) post-DEN. This was compared to tissue constructs combining healthy untreated epithelium and stroma combined (*in vitro* control). Ep, epithelium; Str, stroma; wt, wild-type. Top-right and bottom images show the emerging a tumour-like structure in healthy untreated epithelium, top-down and side-views, respectively. Grafted epithelium expressed nuclear tdTomato in red (nT, from nTnG mice). DAPI, blue; β-catenin, green (CTNNB1), PDGFRα, greyscale; (top right, top-down view; bottom, side-views). Scale bars, 25µm. **b**-**c**, Experimental protocol (left) and representative side-views of 3D-rendered confocal images (right), of 3D heterotypic tissue constructs where both epithelium and stroma are from healthy untreated control animals 7 days post-culture (**b**) and *in vivo* healthy untreated tissue (**c**) for comparison (**b** and **c** are controls for **Fig. 2b-c**). PDGFRα labels fibroblasts (greyscale); in $R26^{Fucci2aR}$ tissue (mCherry, G1 cells; mVenus, S/G2/M cells). Scale bars, 25 µm. **d**, Confocal image of sporadic surviving graft of healthy untreated epithelium nT (red) combined with healthy untreated stroma EGFP (from H2B-EGFP mice). (**d** is control for **Fig. 2e-g**) DAPI, blue. Scale bars, 500µm. Panels (**a**) and (**b**) created with BioRender.com.

#### Extended Data Figure 4

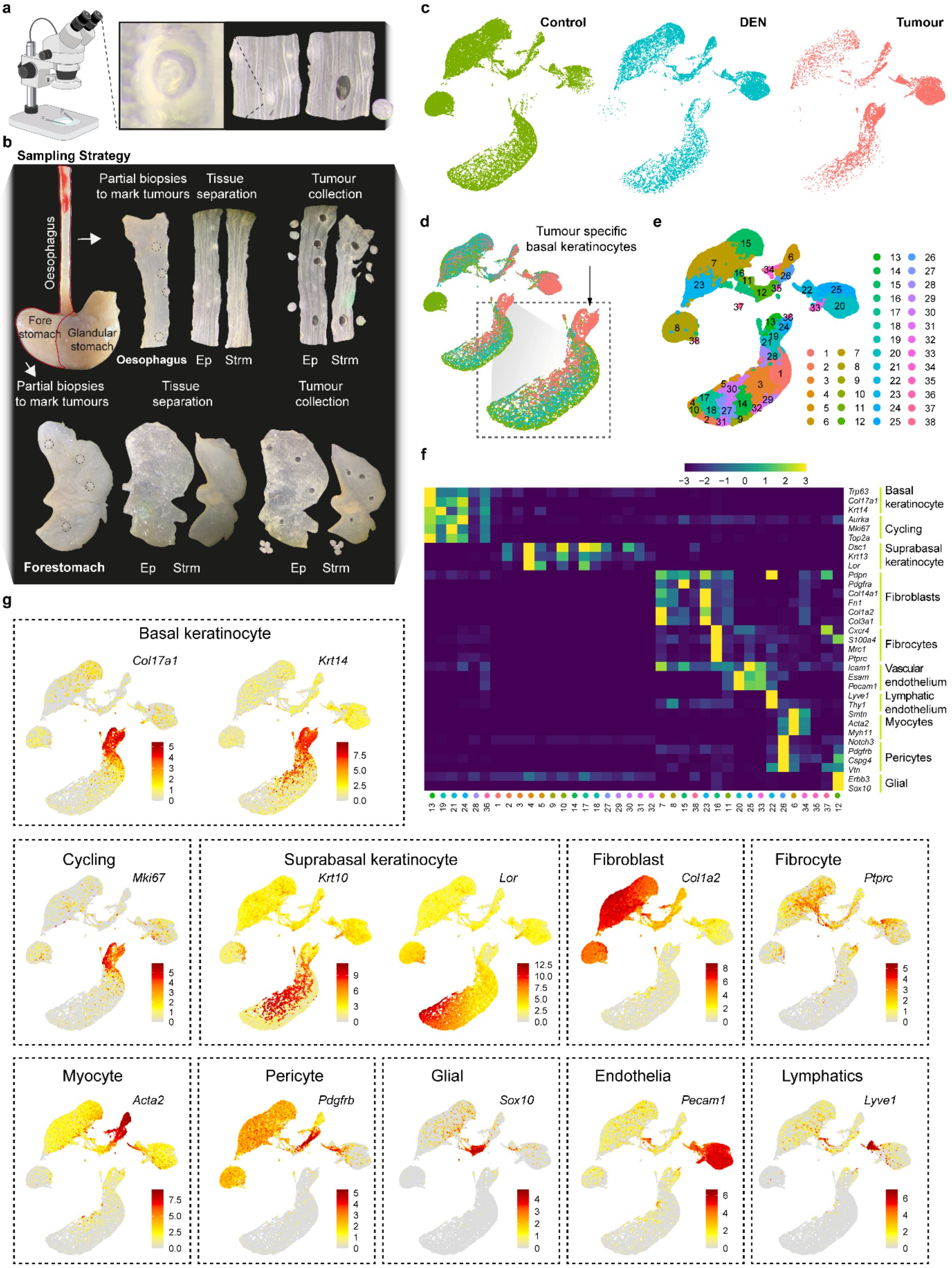

**Extended Data Fig**. **4: Single cell RNA sequencing annotation; (Corresponds to Fig. 3) a,** Schematic showing visualization of representative DEN-induced tumours under the dissection microscope, before and after microdissection using punch biopsy tool. **b**, Image composite depicting sample collection strategy for scRNA-seq. Squamous oesophagus and forestomach (outlined in red) first underwent tumour marking, and were then peeled to separate epithelial (Ep) and Stromal (Str) compartments. This was followed by sample biopsying, where tumours and size-matched control biopsies (adjacent tissue as internal control, DEN; untreated tissue as external control, Control) were microdissected for scRNA-seq. **c**, Cell distribution in the UMAP dimensionality reduction space showing samples of origin for control, DEN, and tumour conditions. **d**, Overlay of UMAP cell distribution from **c**. Inset, highlights area marking tumour specific basal keratinocytes. **e**, UMAP representing cell cluster distribution as defined by Seurat. **f**, Heatmap showing expression of representative marker genes used for cell type annotation detected across the 38 clusters. Log-transformed normalized expression levels were averaged by cluster for each gene and scaled across all cells belonging to each group. Scale bar denotes expression range (scale: -3 to 3). **g**, UMAPs showing expression of representative genes for each cell type identified. Colour bars of UMAPs indicate $\log_2$-transformed normalized expression levels. Panel (**a**) created with BioRender.com.

#### Extended Data Figure 5

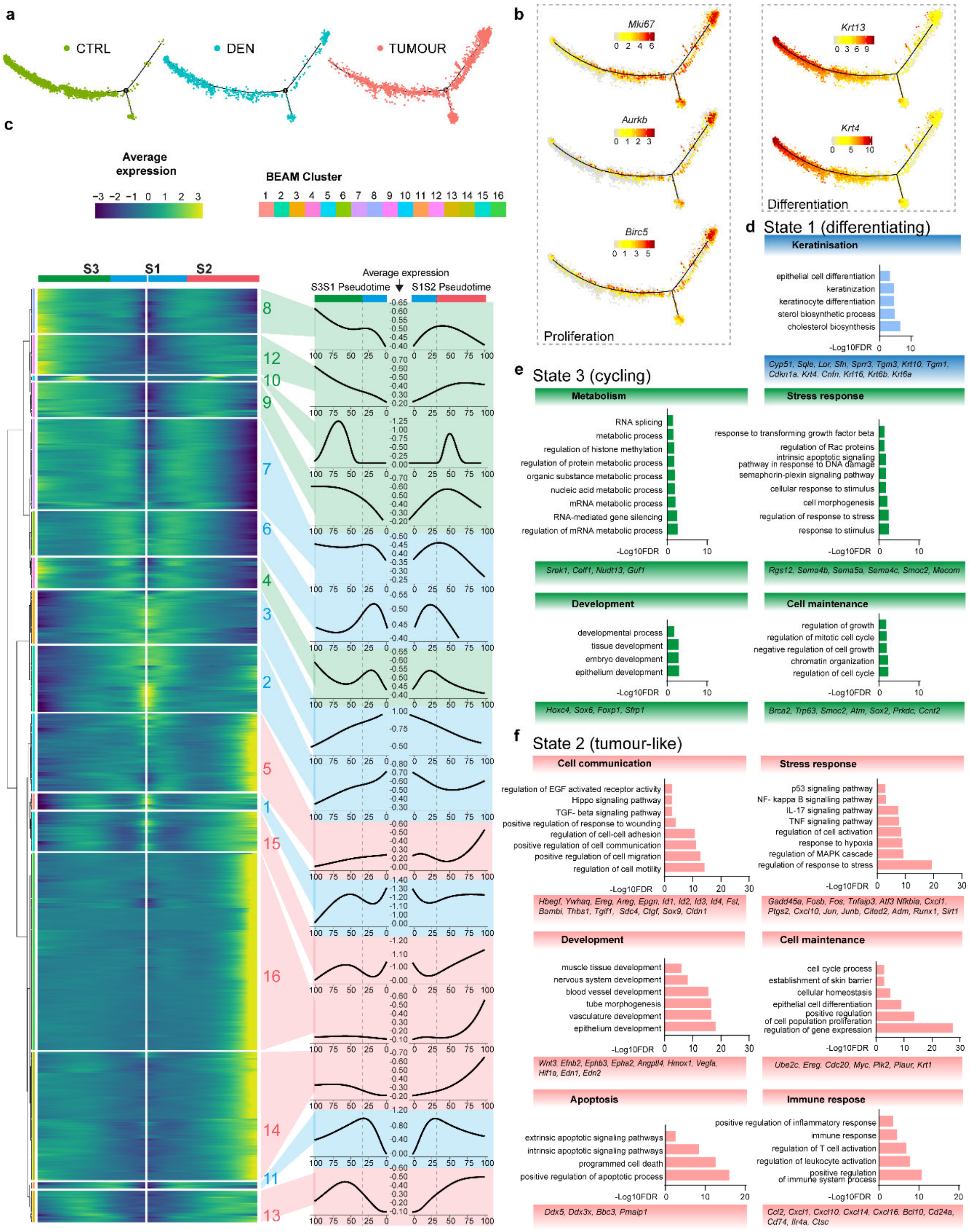

**Extended Data Fig. 5: Pseudotime analysis denotes distinctive early tumour state in the keratinocyte population; (Corresponds to Fig. 3) a**, Basal epithelial cells were ordered along a two-dimensional pseudotime (PST) trajectory. Tumour cells were enriched in state 2. **b**, Expression of proliferation and differentiation maker genes across the PST trajectory. Colour bars indicate $\log_2$-transformed normalized expression levels. **c**, Heatmap (left) of the top 1000 genes identified as differentially expressed between PST states 2 (tumour state) and 3 (basal cycling). $\mathrm{Log}_2$-transformed expression levels were scaled in a gene-wise fashion from -3 to 3 (scale) and are shown after hierarchical clustering along trajectories from S1 to S2 (green to blue) and from S1 to S3 (blue to red). The average expression pattern of the genes contained in each cluster are depicted on the right, with three major pattern types having been identified based on the PST score of peak expression (Blue, peak in state 1; Red, peak in state 2; Green, peak in state 3). Dotted lines label the highest PST score of state 1. **d-f**, Gene enrichment analysis of genes enriched in the different PST states colour coded as per **c**, compiled from Gene Ontology Biological Process (GO:BP) and Reactome (REAC) terms, and grouped based on biological knowledge. Terms with p-value < $1.0\times10^{-16}$ were ranked by their $-\log_{10}$(FDR [false discovery rate]) value. Representative genes are listed below of each plot.

### Extended Data Figure 6

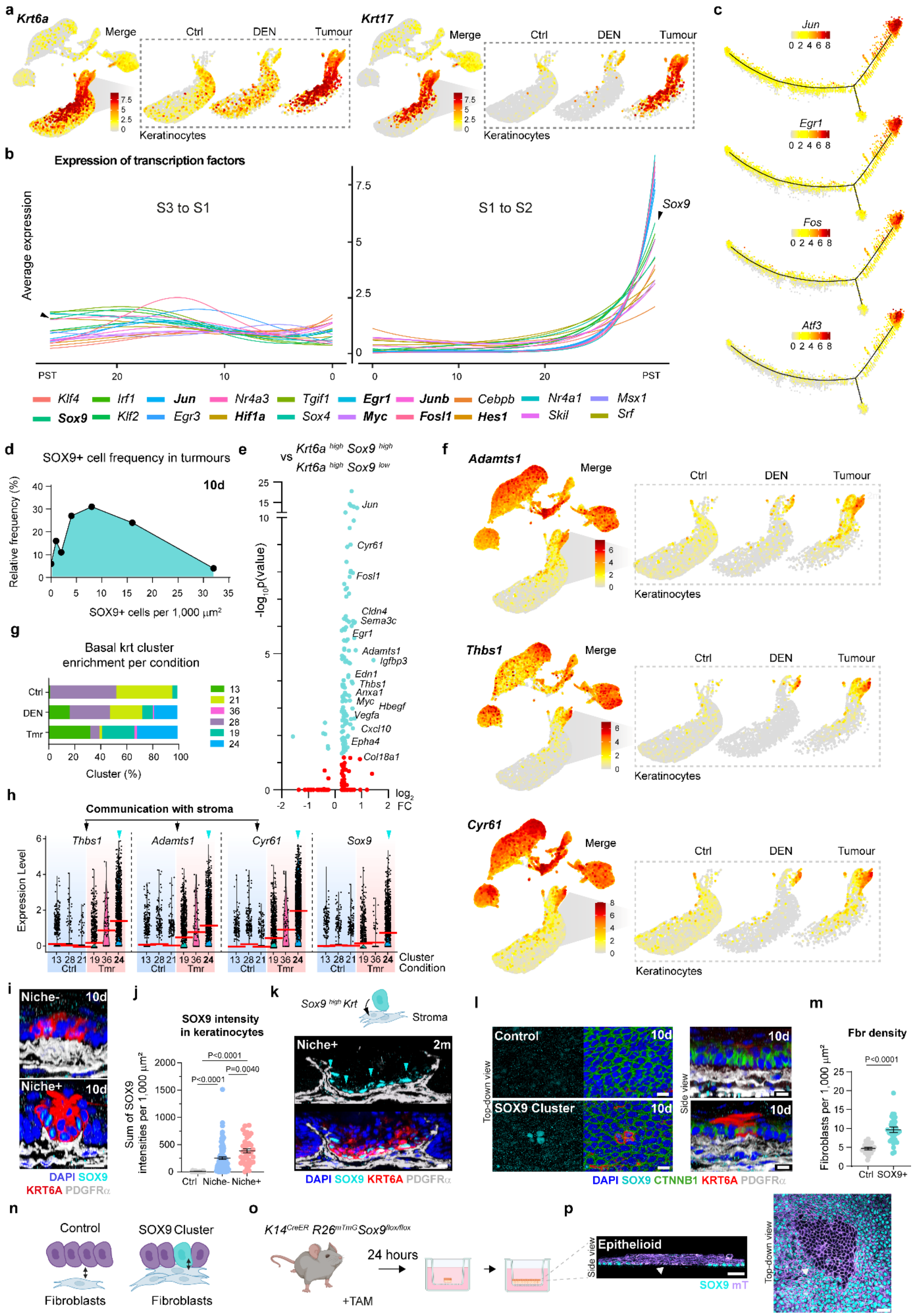

**Extended Data Fig. 6: Transcriptional signature of keratinocyte tumour state denotes features of epithelial-stromal communication. (Corresponds to Fig. 3) a**, UMAPs showing expression of early tumour marker genes. Insets depict keratinocyte clusters split by condition: external control (ctrl), internal control (DEN), and tumours. **b**, Expression of transcription factors along the two pseudotime trajectories: from state (S) 1 to either S3 or S2. Data denote an overexpression of stress-related transcription factors in the tumour state (S2). Gene expression is shown as the number of $log_2$-normalized UMIs detected for each gene across all cells at each of the PST trajectory points after smoothening with the genSmoothCurves function. **c**, Expression of relevant transcription factors plotted in the PST trajectory. Colour bars indicate $log_2$-transformed normalized expression levels **d**, Relative frequency of SOX9+ keratinocytes (%) in tumours 10 days post-DEN, n- number of tumours=119 from 3 biological replicates. **e,** Volcano plot showing differential gene expression between *Krt6a$^{high}$Sox9$^{high}$* and *Krt6a$^{high}$Sox$^{low}$* keratinocytes in PST S2. Non-significant genes ($p \geq 0.05$), red. Genes detected in enriched pathways as identified in **Fig. 3j** are labelled. **f**, UMAPs showing expression of selected genes enriched in **e**; insets depicts keratinocyte clusters split by condition. **g**, Barplot showing the distribution of basal keratinocytes (krt) across clusters split by condition. **h,** Violin plots illustrating $log_2$-transformed normalized expression levels of selected genes in basal keratinocyte clusters split by condition (Ctrl, clusters 13, 28, 21; Tmr, clusters 19, 36, 24). Cyan arrowheads mark the cluster with cells expressing the highest level of *Sox9*. Means denoted by a red line. **i,** 3D-rendered confocal images of niche+ and niche- tumours showing KRT6A (red) and SOX9 (cyan) 10 days (d) post-DEN. Scale bars 10 µm. corresponds to **Fig. 3k,l**. **j,** Sum of intensities of SOX9+ keratinocytes per surface area in niche- or niche+ tumours and control areas. Tumours analysed 10 days post-DEN. n is the number of control areas, niche- or niche+ tumours from 3 mice each; n=19, 84, 35, respectively. One-way Welch ANOVA was used for comparisons. **k,** Cartoon (top) representing SOX9$^{high}$ keratinocyte communication with stroma and a side-view of a tumour 3D rendered confocal image (bottom) 2 months (m) post-DEN. Scale bars, 10 µm. Cyan arrowhead marks SOX9+ keratinocytes in direct contact with the stromal niche. **l,** 3D-rendered confocal images (basal view, left; side-view, right) of SOX9+ keratinocyte clusters exclusively in DEN condition. Scale bar, 10 µm. Corresponds to **Fig. 3m, n. m,** Number of fibroblasts directly underneath the basal layer corrected per area. n is the number of control or SOX9 expressing regions assessed in tissues from 4 mice; n=28, 33, respectively. Two-tailed Welch's t-test. **n,** Cartoon representing the intimate interaction between fibroblasts and SOX9+ keratinocytes in DEN treated tissue compared to control area. **o,** Strategy to validate SOX9 knock-out in basal keratinocytes. SOX9 expression was induced by culturing oesophageal tissue using the 3D epithelioid approach. Established cultures from tamoxifen-induced *Krt14$^{CreER}$ R26$^{mTmG}$Sox9$^{flox/flox}$* mice were confocal imaged in **p**, confirming the presence of areas lacking SOX9 expression. n=3, biological replicates. Scale bar, 25 µm. White arrowheads mark the SOX9 loss. Confocal images **i**, **k**, **l**, **p** show SOX9, cyan; β-catenin (CTNNB1), green; KRT6A, red; membrane Tomato (mT), purple; PDGFRα, greyscale; DAPI, blue. Colour bars of UMAPs (**a** and **f**) indicate $log_2$-transformed normalized expression levels. Panels (**k**), (**n**) and (**o**) created with BioRender.com.

### Extended Data Figure 7

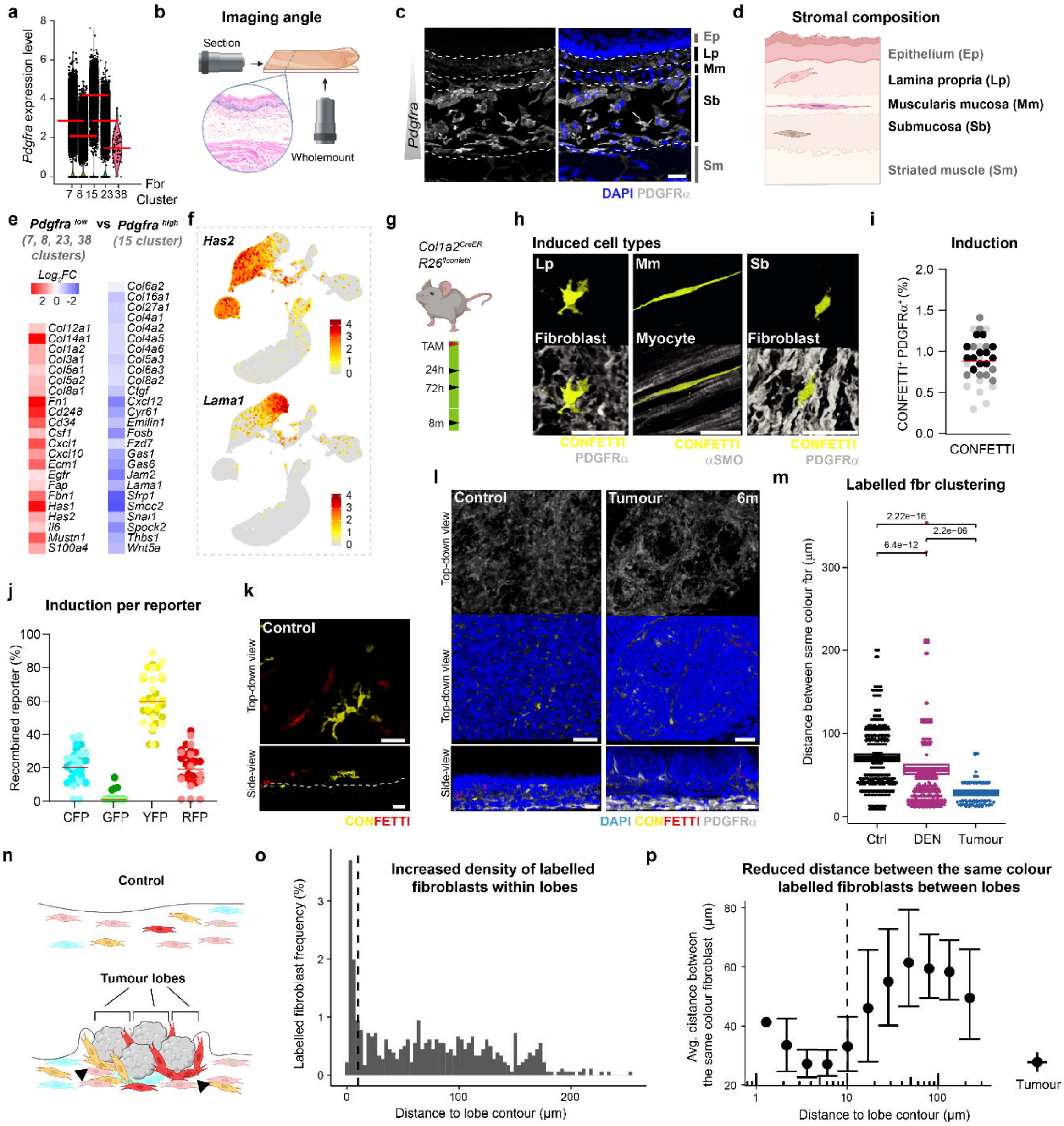

**Extended Data Fig. 7: Expansion of Pdgfra$^{low}$ fibroblast population in the early tumour niche. (Corresponds to Fig. 4) a**, Violin plots showing the levels of *Pdgfra* expression in fibroblast clusters 7, 8, 15, 23, and 38. Mean expression denoted by red line. **b**, Schematic representation of the imaging angle used according to sample preparation method used. **c,** Image of a tissue section showing reduced PDGFRα expression in upper stromal layers (i.e. lamina propria). Dashed lines label tissue layers as listed in schematic **d**, showing tissue layers of upper gastrointestinal tract. **e,** Heatmap showing differentially expressed genes between *Pdgfra$^{low}$* and *Pdgfra$^{high}$* fibroblast populations. $Log_2$-transformed normalized expression levels were averaged by clusters for each gene. Scale bar denotes $log_2$-transformed normalized expression levels. **f**, UMAPs showing the expression levels of genes characterising the *Pdgfra$^{low}$* (*Has2*) and *Pdgfra$^{high}$* (*Lama1*) populations. Color bars indicate $log_2$-transformed normalized expression levels **g,** Experimental protocol of fibroblast lineage tracing in *Col1a2$^{CreER}$Rs26$^{FConfetti}$* mice; hours (h), months (m). **h**, Confocal images of confetti-labelled collagen-expressing cells 24-hours after induction in different stromal layers. Scale bars, 25 µm. **i-j**, Efficiency of confetti construct recombination after a 72-hour chase in the PDGFRα+ fibroblast population. Total percentage of recombined cells in the PDGFRα+ fibroblast population (**i**), and split colour percentage within the recombined population (**j**). n represents number of fields from 3 animals, n= 36. **k-l**, Confocal 3D top-down or side-view images of a control area or a tumour 6 months (m) post-DEN. Corresponds to **Fig. 4c,d**. Dashed line marks the surface of the submucosa (in **k**). **m,** Distance to the nearest neighbour fibroblast (fbr) labelled in the same colour in external control (Ctrl), internal control (DEN) and tumours. The horizontal line denotes the average, and the box signifies 95% confidence intervals obtained via bootstrapping. P-values were obtained using a Wilcoxon test. **n**, Cartoon illustrating fibroblast expansion in the early tumour niche: lineage tracing results revealed that stroma between tumour lobes show increased fibroblast density and decreased distance between the same colour fibroblasts (black arrowheads) if compared to the tissue outside of lobes or control. **o,** Histogram showing the percentage of cells in a given bin of the distance to the nearest lobe contour. **p,** The average distance to the nearest neighbour of the same colour as a function of the distance to the lobe boundary. Results show that labelled cells are closer together on the lobe contour than elsewhere. The vertical dashed line in **o** and **p** denote 10 µm, the estimated width of the tumour lobe contour. Confocal images in **c**, **h**, **k**, **l** show PDGFRα, greyscale; Confetti, yellow and red; PDGFRα and αSMO, greyscale. Panels (**b**), (**d**), (**g**) and (**n**) created with BioRender.com.

#### Extended Data Figure 8

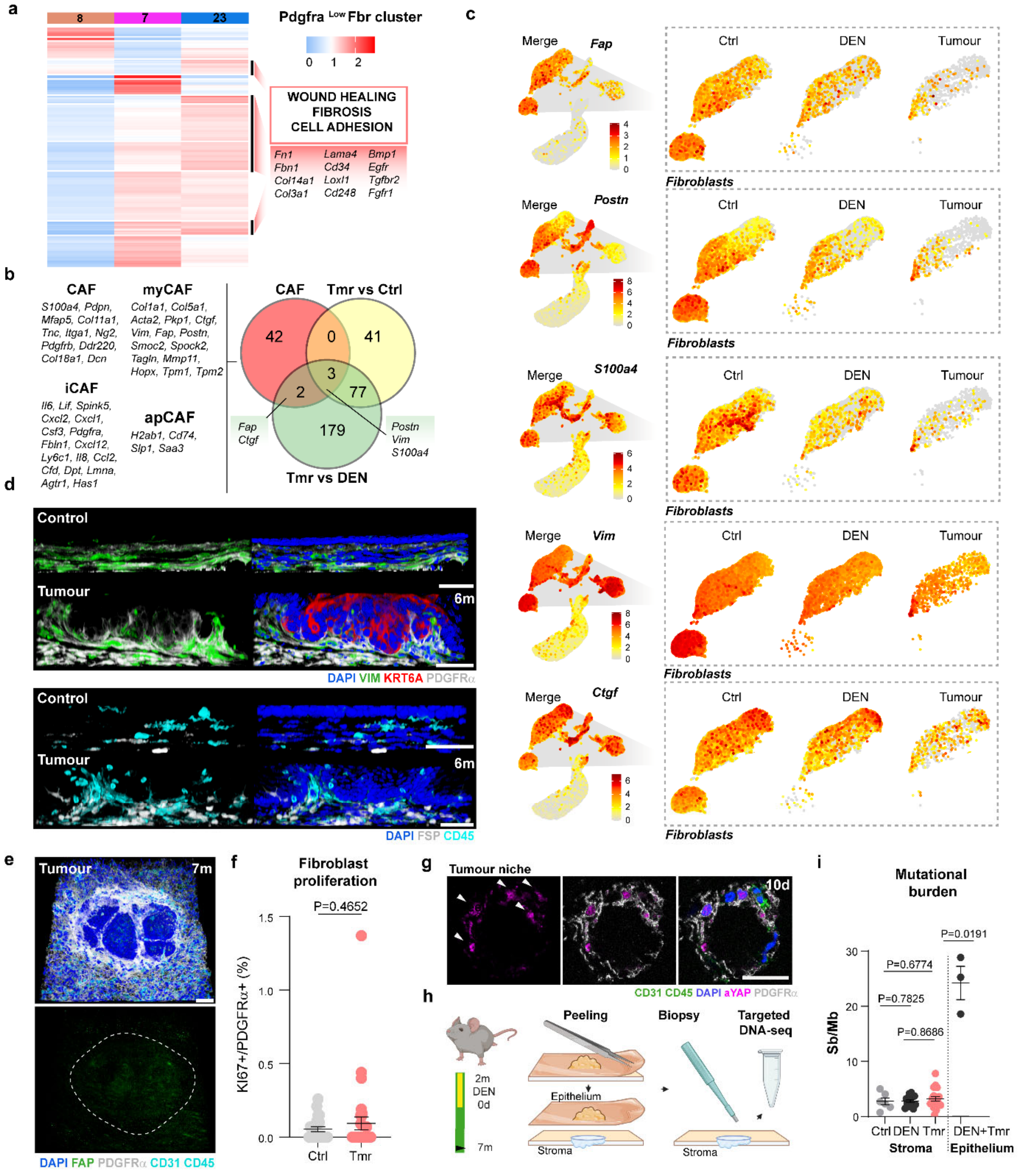

**Extended Data Fig. 8: Tumour niche fibroblasts transition towards Cancer Associated Fibroblast (CAF) identity. (Corresponds to Fig. 4) a**, Heatmap (left) showing differentially expressed genes ($|Log_2FC| > 0.5$, $p < 0.05$) in Cluster 23 compared to the remaining *Pdgfra$^{low}$* fibroblast clusters. $Log_2$-transformed normalized expression levels were averaged by cluster for each gene and scaled across all cells belonging to each group. On the right, terms enriched in Cluster 23 selected based on GO:BP terms and literature mining. Representative genes are listed below. **b**, Venn diagram displaying overlapping genes between known CAF markers (top left, red) and differentially expressed genes (DEGs) in DEN tumour (Tmr) fibroblasts (Cluster 23) relative to external control (Ctrl; top right, yellow) or to internal control cells (DEN; bottom, green). Overlapping genes are listed. Known subtypes of CAFs: Myofibrotic (my), immune (i), and antigen-presenting (ap) were considered. **c**, UMAPs showing expression of signature genes found to be enriched in Cluster 23 tumour fibroblasts; insets display fibroblast clusters split by condition: external control (ctrl), internal control (DEN), and tumours. Color bars indicate log2-transformed normalized expression levels. **d**, Representative 3D-rendered side-view images of control tissue and tumours 6 months (m) post-DEN. Experession of CAF markers was not detected in niche fibroblasts. KRT6A, red; CD45 (immune), cyan; Vimentin (VIM), green; PDGFR$\alpha$ (top) and FSP (bottom), greyscale; DAPI, blue. Scale bars, 50µm. **e**, 3D-rendered confocal image of tumour stroma 7 (m) months post-DEN. Dashed line outlines tumour area. Fibroblast activated protein (FAP), green; PDGFR$\alpha$, greyscale; CD31 and CD45, cyan; DAPI, blue. Scale bars, 50 µm. **f**, Quantification of Ki67+ EGFP+ fibroblasts from *Pdgfra$^{EGFP}$* mice in control and niche+ tumours 6-12 months (m) post-DEN. n, control areas or tumours, n=25, 50, respectively, from 7-8 animals. Data is expressed as mean ± s.e.m. Two-tailed Mann Whitney test. **g**, Confocal images of tumour stroma 10-days post-DEN. Images show active (a) YAP expression in PDGFR$\alpha$ fibroblasts (white arrowheads). aYAP, magenta; CD31 and CD45, green; PDGFR$\alpha$, greyscale; DAPI, blue. Scale bars, 25µm. **h,** Experimental protocol for targeted DNA sequencing of stromal niche. **i**, Substitutions per megabase (Sb/Mb) were calculated and used as indicative of the mutation burden in stroma across conditions. n is the number of tissue biopsies or tumours analysed, n=6, 13, 18, respectively. The epithelial tissue (12 months post-DEN treatment) was used as benchmark and includes both tumour and tumour-free tissues (DEN+Tmr). For the epithelium, whole oesophagi from 3 mice were used. Data is expressed as mean ± s.e.m. Two-tailed Welch's *t*-test comparing tumour stroma with internal or external control stromas or with epithelium. Panel (**h**) created with BioRender.com.

#### Extended Data Figure 9

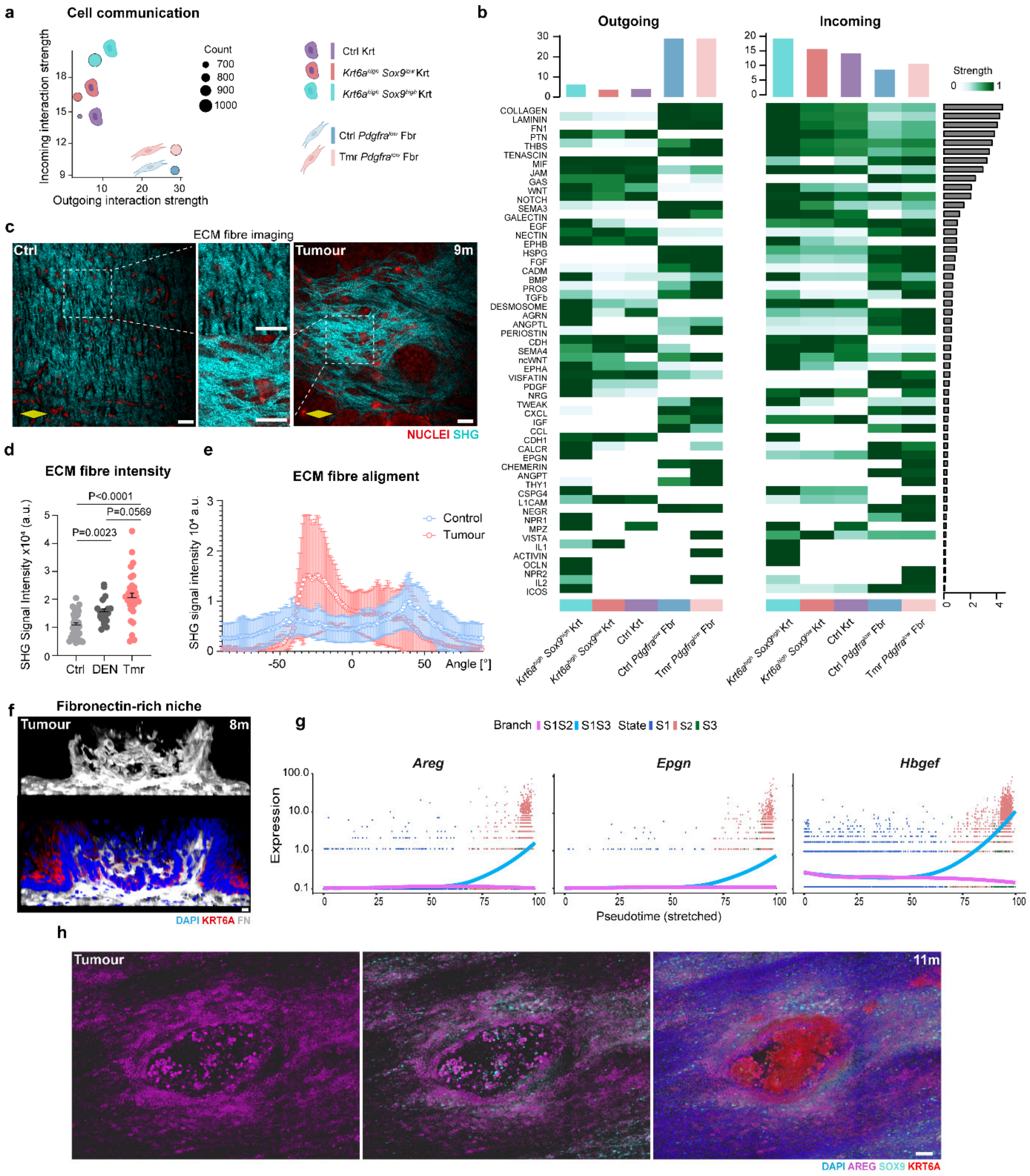

**Extended Data Fig. 9: The early tumour niche is characterised by a pro-fibrotic environment (Corresponds to Fig. 5) a**, Scatterplot displaying the dominant senders and receivers identified as predicted by CellChat. Circle size represents 'communication probabilities'. Illustrations of cell types are as in **Fig 5a**-**b**, Heatmaps representing outgoing and incoming interaction strengths rescaled to their row maxima. Cumulative (total) interaction strength is depicted by bars on the right. Bar plots above indicate total interactions per cell type. Control, ctrl; Tumour, Tmr; Keratinocyte, Krt; Fibroblast, Fbr. **c**, Second harmonic generation (SHG) image of lamina propria (directly underneath epithelium) in a control (Ctrl) and tumour areas 3 months (m) post-DEN. Dashed lines mark insets. Extracellular matrix (ECM) fibres detected by SHG, cyan; nuclei, red. Scale bars, 25 µm. The yellow diamonds represent the longitudinal orientation of the oesophagus **d**, ECM fibre density was scored as SHG signal intensity 3 months post-DEN. n is the number of regions or tumours assessed from 5-6 animals; n=36 (Ctrl), 20 (DEN), 31 (Tmr), respectively. Bars indicate mean + s.e.m. One-way Welch's ANOVA. **e,** Plot depicting the orientation of ECM fibres in control regions and tumours 9 months post-DEN. n is the number of fields of view assessed in 3 animals per condition, n=12 (Ctrl) and 8 (Tmr). **f,** Confocal 3D-rendered side-view of a tumour 8 months post-DEN. KRT6A, red; Fibronectin (FN), greyscale; DAPI, blue. Scale bar, 25 µm. **g,** Two divergent kinetic trends along the PST trajectories from S1 to S2 (S1S2) and from S1 to S3 (S1S3) show upregulation of the EGF ligands *Areg*, *Epgn* and *Hbegf* in S2 (tumour state). **h**, Confocal images of tumour epithelium 11 months post-DEN confirms expression upregulation of EGFR ligand AREG. AREG, magenta; SOX9, cyan; KRT6A, red; DAPI, blue. Scale bar, 50 µm.

### Supplementary Methods

#### Experimental mouse lines

To assess the proliferative state of cells, *R26*$^{Fucci2aR}$ (*Fucci2a*) mice were used. This mouse line constitutively expresses fluorescent reporters that highlight different phases of the cell cycle (G1 is marked by *mCherry-hCdt1*, and S/G2/M by *mVenus-hGem*). To identify fibroblasts, *Pdgfra*$^{EGFP}$ mice were used. These mice constitutively express an H2B-eGFP fusion peptide from the endogenous *Pdgfra* locus. For *Sox9* conditional knock-out experiments, *Sox9*$^{flox/flox}$ were crossed with *K14-Cre*$^{ER}$ mice to generate *K14*$^{CreER}$/ *Sox9*$^{flox/flox}$ animals in which *Sox9* expression is inactivated in recombined epithelial cells upon tamoxifen (TAM) administration. To visualise cells in 3D cultures (epithelioid)[1] cells *in vitro* the fluorescent reporter mouse line *R26*$^{mT-mG}$ (*mTmG*), which constitutively express tdTomato localised in the cell membrane, was crossed with *K14*$^{CreER}$/*Sox9*$^{flox/flox}$ and the conditional knock out phenotype was visualised *in vitro*. For fibroblast lineage tracing, *Col1a2*$^{CreER}$ mice expressing a Tamoxifen-inducible *Cre* recombinase under the control of the endogenous *Col1a2* promoter were used. This line was crossed with *R26*$^{FlConfetti}$ mice in which the Rosa26 locus was targeted with a transgenic cassette containing the sequences for stochastic expression of either cyan, green, yellow or red fluorescent protein (CFP, GFP, YFP and RFP) upon *Cre* recombination. Thus, by using low doses of TAM in *Col1a2*$^{CreER}$/ *R26*$^{FlConfetit/WT}$, individual Col1a2-expressing cells and their progeny can be labelled with one of these four fluorescent reporters. To identify the origin of tissues in 3D tissue recombination culture assays (donor or recipient), *R26*$^{nT-nG}$ (*nTnG*) and *H2B-EGFP* (*CAG::H2B-EGFP*) mice were used. *R26*$^{nT-nG}$ ubiquitously express a *tdTomato* fluorescent reporter that localises to the nucleus, while *H2B-EGFP* mice constitutively express EGFP that localizes to the nucleus. In transplantation assays, mice commonly known as NOD scid gamma (NSG; NOD.Cg-*Prkdc*$^{scid}$ *Il2rg*$^{tm1Wjl}$/SzJ) were used as recipients. Due to the absence of *Prkdc* and X-linked *Il2rg* expression, these mice are immunodeficient.

#### Analysis of migration assay

The total number of fibroblast and the number of fibroblasts that crossed the transwell insert membrane were counted and migration rates calculated as the fraction of migrated fibroblasts relative to all fibroblasts detected. Membranes from transwell inserts were visualised by their autofluorescence after excitation with the 405 laser. Displayed images were produced using Volocity 5.3.3.

#### Assessment of tumour phenotype, size and number in whole-mounts

To identify tumours in the upper gastrointestinal tract as early as 10 days after DEN withdrawal, we used DAPI to asses changes in tissue topology, as well as cell morphology. This was combined with immunostaining against Keratin 6A (KRT6A) - a tumour-marking keratin[2]. PDGFRα was used as a marker of residential fibroblasts to asses niche phenotype. Approximately one third of the oesophagus (middle part) was imaged using the following settings: a 40× objective, zoom 0.75, an optimal pinhole size (as defined by the software), a scan speed of 400 Hz, a line average of 1, a Z-step size of 1.5 μm, and a resolution of 512 × 512 pixels. Tumour phenotype (niche- or niche+) was visually scored by assessing basal keratinocyte morphology, the level of fibroblast recruitment, and fibroblast remodelling in a single-tile section view using Volocity 5.3.3. Tumour diameter was scored in Volocity by measuring the distance between the first and last tumour nuclei along the transversal tumour axis in one plane. The number of tumours detected per tissue was normalised to the surface area analysed (typically 10-50 $mm^2$). To estimate the number of tumours per oesophagus, the surface areas of 4 whole oesophagi were measured and the total number of tumours extrapolated accordingly.

**Proliferation, SOX9 and stromal cell distribution measurements in whole-mounts**

EdU$^+$ and SOX9$^+$ tumour keratinocytes were quantified by manually cropping the stromal part of 3D images and running an automated pipeline for nuclear signal detection in Volocity 5.3.3. Control measurements were created by cropping regions of equivalent size in control areas. The number of identified objects or the sum of mean object signal intensities were normalised to surface area of each tumour, which was calculated by using the formula $S= \pi ab$ (a, width; b, height). The latter assumes that, in a single XY plane, the shape of an oesophageal tumour is an ellipse.

SOX9 mean intensities per object were summed per area of interest and normalised to the surface area of interest as well as to the individual sample average (self-average) to account for differences in intensity across imaging sessions.

Stromal cell distribution in niche+ and niche- tumours was estimated by counting all stromal cells in the lamina propria directly underneath the tumour epithelium and subsequently normalising this number to the tumour XY cross-sectional area. Cells were then identified as fibroblasts (PDGFRα$^+$), immune (CD45$^+$) or endothelial (CD31$^+$) and their numbers expressed as a percentage of total stromal cells.

Fibroblast proliferation in tumours was scored by manually counting Ki67$^+$ EGFP$^+$ cells in *Pdgfra*$^{EGFP}$ DEN-treated mice. The number of proliferating fibroblasts was normalised to the total number of fibroblasts in the lamina propria directly underneath the epithelial tumour.

**Fibroblast density and proximity to SOX9$^+$ keratinocytes**

Oesophageal tissues collected 10 days after DEN withdrawal were used to source SOX9$^+$ keratinocytes outside the tumour area. These were KRT6A-expressing, SOX9-expressing cell clusters (between 5-28 cells in size) that had normal topology as assessed by the position of DAPI-labelled nuclei. Tissues were additionally immunostained with antibodies against β-catenin to identify keratinocytes and against PDGFRα to identify fibroblasts. The number of fibroblasts in the lamina propria region directly underneath SOX9$^+$ keratinocyte clusters was counted manually and normalised to the surface area considered; equivalently sized areas in tissues of untreated animals were used as controls. The distance between SOX9-labelled keratinocyte clusters and the 5 closest fibroblasts was manually measured in either an xz or an yz cross-section view using Volocity 5.3.3 and is expressed as an inverse distance to demonstrate proximity.

**Fibroblast lineage tracing**

Oesophagi from *Col1a2*$^{CreER/WT}$ *R26*$^{FlConfetti/WT}$ mice that received TAM at the beginning of the DEN treatment were collected 6 months after DEN withdrawal. Tissues were processed and stained for PDGFRα and DAPI as described above. The more abundant YFP$^{+}$ and RFP$^{+}$ clones within tumours and in normal tissue regions were imaged on a confocal microscope and analysed with Volocity 5.3.3. To investigate clonal expansion of YFP$^{+}$ and RFP$^{+}$ fibroblasts, the coordinates of the tumour lobe (defined as the region between its basal keratinocyte layer and the adjacent lamina propria) were recorded in XY at intervals of 1-2 µm; the process was repeated at different Z positions throughout the entirety of the tumour (every 1-4 µm). Similarly, the XYZ coordinates of the nucleus of each confetti-labelled fibroblast found within the field of view as well as its colour were recorded. The average distance to the nearest fibroblast neighbour of the same colour was calculated and normalised to the distance to the nearest fibroblast neighbour of a different colour. Statistical significance was assessed using the Mann-Whitney U test (Wilcoxon signed-rank test).

**Second-harmonic generation (SHG) image analysis**

Using the Fiji software for bioimage analysis, a region of 200x200 pixels was arbitrarily selected within the lamina propria in each control image. In the case of tumour images, such region was selected to be directly underneath the lesion site. Fibre alignment was measured within these regions using the OrientationJ plug-in for Fiji[3] with the σ tensor value set to 7. For determining the emission intensity of extracellular matrix fibrils, images of the lamina propria were first exported in the TIFF format using ZEN. Within each control image, intensity was measured in four equivalent circular regions (phantom tumours; diameter = 125 pixels) and averaged. Only one such region immediately under the lesion site was considered for tumour images.

**Differential gene expression and gene set enrichment analysis (GSEA)**

Based on the expression dynamics of the genes in each of the branched heatmap clusters throughout pseudotime, clusters were grouped (colour-coded in **Extended Data Fig. 5c**) and the genes within them used for Gene Ontology analysis with g:profiler (https://biit.cs.ut.ee/gprofiler/gost version e109_eg56_p17_1d3191d) and STRING (https://string-db.org/ version 12.0). Gene Ontology biological processes (GO:BP) were grouped to mother terms in Revigo (http://revigo.irb.hr/ version 1.8.1). The resulting GO:BP mother terms, Kyoto Encyclopaedia of Genes and Genomes (KEGG) (https://doi.org/10.1093/nar/28.1.27) terms and Reactome (REAC) (https://doi.org/10.1093/database/baz123) terms were manually grouped, with the most representative significant terms selected for depiction based on experimental observations and biological knowledge.

By focusing on the clusters from the heatmap generated by the plot_genes_branched_heatmap function (**Extended Data Fig. 5c**) that contain genes displaying an increased expression towards the end of the S1 to S2 trajectory, we identified 49 previously described murine transcription factors (TFs). From those, the expression dynamics of the 20 with the most striking increase was extracted across each of the pseudotime trajectories (S1S2 and S1S3; **Extended Data Fig. 6b**) using the genSmoothCurves function as described under the "**Cell transition trajectory analysis**" section above using standard parameter settings (except for the relative_expr parameter which was set to FALSE). The values generated were subsequently normalized to the average expression of each gene throughout the specific PST trajectory (S1S2 and S1S3) and were plotted as overlaid curves using the R package ggplot2 (v3.5.0) (**Extended Data Fig. 6b**).

Comparing the gene expression profile of *Krt6a*$^{high}$*Sox9*$^{high}$ keratinocytes against that of *Krt6a*$^{high}$*Sox9*$^{low}$ ones found in PST state 2 (642 and 655 cells, respectively), allowed us to detect 276 DEGs. Among them, 119 had a $\log_2$ fold-change > 0.25 and a p-value < 0.05. Using these 119 genes, we conducted GSEA as described above. Most of the relevant terms identified are depicted in **Fig. 3j**.

The expression of three genes, namely *Areg, Hbegf* and *Epgn,* was plotted throughout PST with separate but overlaid curves for each of the two detected trajectories (S1S2 and S1S3) using the monocle function plot_genes_branched_pseudotime with standard parameter settings, except for the relative_expr parameter which was set to FALSE. Importantly, values on the x-axis are shown for both trajectories on an adjusted scale of 0 to 100, which represent 100 evenly spaced PST time points.

Fibroblasts expressing *Pdgfra* were categorised as either *Pdgfra*$^{low}$ (clusters 7, 8, 23 and 38) or *Pdgfra*$^{high}$ (cluster 15) and genes differentially expressed between them were identified using the FindMarkers function of the Seurat package with standard parameters. The $\log_2$ fold-change of significantly regulated genes encoding proteins contained in the matrisomereferen was plotted as a heatmap using GraphPad Prism.

*Pdgfra*$^{low}$ fibroblast sub-types were additionally compared against each other. The average expression per cluster for DEGs identified from comparing the fibroblasts in Cluster 23 against the rest of *Pdgfra*-low fibroblasts (Clusters 7, 8 and 38 all together) was extracted for each cluster (except for clusters 8 and 38, which were merged into a single cluster [8+38] due to the low number of cells in the latter). Values were subsequently normalised so that the average expression of each gene across all clusters (7, 23, 8+38) was assigned a value of 1. Normalised values were subsequently plotted as a heatmap (**Extended Data Fig. 8a**).

Similarly, in cluster 23, DEGs between conditions (namely, Tumour vs control and Tumour vs DEN) were identified using the FindMarkers function of the Seurat package. DEGs with a p-value < 0.05 and a $\log_2$ fold-change above 0.5 were cross-referenced with known cancer-associated fibroblast (CAF) markers. The overlap is schematically shown in a Venn diagram (generated in the website https://www.bioinformatics.org/gvenn/)[4,5].

**Supplementary Table 1.** Antibodies list

| Antibody | Dilution | Company | Cat number | |
|---|---|---|---|---|
| AREG | 1 in 50 | R&D Systems, Inc | AF989 | Primary |
| aSMA | 1 in 200 | Abcam | ab5694 | |
| β-Catenin | 1 in 500 | Cell Signaling Technolo | 9562 | |
| CD31 | 1 in 200 | Abcam | ab7388 | |
| CD45 | 1 in 200 | BioLegend | 103102 | |
| FAP | 1 in 200 | Abcam | ab218164 | |
| FN | 1 in 200 | BD Biosciences | 610078 | |
| K17 | 1 in 200 | Proteintech | CL488-17516 | |
| KRT6A | 1 in 1000 | BioLegend | 905701 | |
| KI67 | 1 in 500 | Abcam | ab16667 | |
| PDGFRA | 1 in 200 | R&D Systems, Inc | AF1062 | |
| S100A4 (FSP) | 1 in 200 | Fisher Scientific Ltd | PA5-16586 | |
| SOX9 | 1 in 1000 | Millipore | AB5535 | |
| VIM | 1 in 200 | Abcam | ab194719 | |
| active YAP | 1 in 250 | Abcam | ab205270 | |
| Goat IgG 647 | 1 in 500 | Millipore | AP180SA6 | Secondary |
| Goat IgG 750 | | Abcam | ab175745 | |
| Mouse IgG 488 | | Invitrogen | A-21202 | |
| Mouse IgG 647 | | Invitrogen | A-31571 | |
| Mouse IgG 750 | | Abcam | ab175738 | |
| Rabbit IgG 488 | | Invitrogen | A-21206 | |
| Rabbit IgG 555 | | Invitrogen | A-31572 | |
| Rabbit IgG 647 | | Invitrogen | A-31573 | |
| Rabbit IgG 750 | | Abcam | ab175728 | |
| Rabbit IgG | | Abcam | ab171870 | |
| Rat IgG 647 | | Abcam | ab150155 | |
| Rat IgG 750 | | Abcam | ab175750 | |
